# Temporal dynamics and functional divergence of the chloroplast division apparatus in *Oryza sativa*

**DOI:** 10.64898/2026.06.26.734771

**Authors:** Caijin Chen, Lei Hua, Andrew R. G. Plackett, Ana Rita Borba, Na Wang, Ruth M. Donald, Susan Stanley, Kumari Billakurthi, Tina Schreier, Tianshu Sun, Julian M. Hibberd

## Abstract

- Chloroplast division is governed by a conserved protein machinery, yet empirical characterization of these regulators remains limited in rice, a primary target for C_4_ engineering. Increased chloroplast occupancy in bundle sheath cells is a hallmark of the C_4_ pathway and so manipulating division is a potential strategy to achieve this goal.
- Through developmental transcript profiling and image analysis, we identified a discrete window of active chloroplast proliferation in rice leaves, coinciding with peak expression of conserved plastid division genes. Functional characterization via overexpression revealed regulatory behaviours distinct from those in *Arabidopsis thaliana*. Overexpression of *OsFtsZ1&2* resulted in fewer, enlarged chloroplasts per bundle sheath cell, whereas *OsMCD1&OsMinE* restricted plastid expansion without altering division rates. Conversely, overexpressing *OsPDV1&2* or *OsARC6&OsDRP5B* increased plastid size without affecting total count. When Os*PDV1&2* were co-expressed with transcriptional regulator *ZmG2*, we observed modest increases in chloroplast size alongside reduced stomatal aperture, increased stomatal density, and higher intrinsic water-use efficiency.
- The results define the temporal landscape of plastid biogenesis in rice and demonstrate divergence across lineages. Our findings suggest that manipulating the division apparatus is insufficient to drive C_4_-like chloroplast biogenesis in the rice bundle sheath, highlighting the complexity of plastid-host cell coordination in cereals.

## Introduction

Meeting rising global food demand amid increasing climatic variability is a central challenge for agricultural sustainability. Enhancing photosynthetic efficiency is a promising route to improve crop yields, given its role in biomass production and untapped genetic potential (Sage & Zhu, 2011; Long *et al*., 2015; Long *et al*., 2025). Strategies range from optimizing light harvesting (Ort *et al*., 2015) to accelerating CO_2_ fixation via enzyme engineering (Croce *et al*., 2024). Among these, installing C_4_ photosynthesis into C_3_ crops would be particularly transformative, with the potential to increase yields by 50% while simultaneously improving water and nitrogen use efficiencies (Hibberd *et al*., 2008). C_4_ species account for nearly 25% of global primary productivity despite comprising only ∼3% of flowering plants - a testament to their high photosynthetic performance (Sage & Zhu, 2011). This advantage arises from coordinated leaf anatomy modifications, cell-type-specific metabolic partitioning, and robust chloroplast biogenesis in the bundle sheath, which together establish a carbon-concentrating mechanism that suppresses photorespiration (Hatch, 1987)

A critical barrier to C_4_ engineering is the paucity of chloroplasts in C_3_ bundle sheath cells. In rice, chloroplasts occupy only ∼14% of bundle sheath cell volume (Karki *et al*., 2013; Wang *et al*., 2017; Lambret Frotte *et al*., 2025), compared to 60–70% in typical C_4_ leaves (Lee *et al*., 2023). While overexpressing transcription factors such as *G2*, *CGA1*, *BZR1*, and *HAP3* can enhance bundle sheath chloroplast development by approximately 30%, these gains remain insufficient to support a functional C_4_ cycle (Hudson *et al*., 2013; Wang *et al*., 2017; Lee *et al*., 2021; Cackett *et al*., 2025; Lambret Frotte *et al*., 2025). Because chloroplast occupancy is a function of both organelle size and number (Cackett *et al*., 2022), and previous efforts have primarily influenced size, we hypothesized that direct manipulation of the chloroplast division machinery could provide a synergistic route to increase bundle sheath chloroplast abundance.

Chloroplast division is mediated by a conserved protein complex spanning the inner and outer envelope membranes. The stromal FtsZ1 and FtsZ2 proteins form a Z-ring at the division site (Vitha *et al*., 2001), positioned by Min proteins (ARC3, MinD, MinE, and MCD1) (Colletti *et al*., 2000; Nakanishi *et al*., 2009; Chen *et al*., 2018). This ring is anchored to the inner membrane by ARC6 and PARC6 (Glynn *et al*., 2008; Glynn *et al*., 2009), which then recruit outer envelope proteins PDV1 and PDV2 to engage the dynamin-related protein DRP5B for final constriction (Miyagishima *et al*., 2006). Studies in *Arabidopsis thaliana* and bryophytes demonstrate that these components are dosage-sensitive. For instance, overexpressing *FtsZ1/2* or *MinE* typically inhibits division, leading to fewer but larger chloroplasts (Colletti *et al*., 2000; Stokes *et al*., 2000; Itoh *et al*., 2001; Schmitz *et al*., 2009), while *PDV1/2* overexpression can increase plastid number (Okazaki *et al*., 2009). However, the functional roles of these genes in monocotyledons remain largely unresolved. While cereal studies have correlated plastid proliferation with leaf developmental gradients (Loudya *et al*., 2021), the individual contributions of division genes to size and number have not been systematically tested in crops. This knowledge gap limits our ability to rationally engineer chloroplast occupancy in the rice bundle sheath.

Here, we investigate whether mis-expression of plastid division genes in rice produces phenotypic effects analogous to those in *Arabidopsis*, and whether combining these with *ZmG2* overexpression yields additive effects on BS chloroplast occupancy. We show that while *OsFtsZ1&2* overexpression disrupts division similarly to eudicot models, *OsMCD1* & *OsMinE* unexpectedly restricted plastid expansion rather than inhibiting division. Furthermore, overexpressing *OsPDV1&2* and *OsARC6&OsDRP5B* enlarged chloroplasts without altering their total number. Notably, co-expression of *ZmG2* with *OsPDV1&2* altered stomatal development, resulting in smaller, more numerous guard cells. This modification decreased stomatal conductance while maintaining CO_2_ assimilation rates, thereby enhancing intrinsic water-use efficiency. These findings reveal distinct regulatory behaviours in rice compared with those previously reported from other species, and indicate that manipulating the division apparatus may offer a tool for improving physiological traits such as water use in crops.

## Materials and Methods

### Plant and growth conditions

Seeds of *Oryza sativa* ssp. japonica cv. Kitaake (wild type) were surface-sterilized, imbibed in sterile Milli-Q water at 30 °C in the dark for 2 d, and germinated on moistened Whatman filter paper in a growth cabinet at 28 °C under a 16/8 h light/dark cycle. After 2 d, seedlings were transplanted to 9 × 9 cm pots (two plants per pot) filled with Profile Field and Fairway soil amendment (Rigby Taylor, UK). Plants were grown in a walk-in growth chamber under a 12-h photoperiod (28 °C day, 20 °C night, with a photon flux density of 400 µmol m⁻² s⁻¹). Plants were fertilized with Peters Excel Cal-Mag Grower solution (LBS Horticulture, UK) supplemented with chelated iron (Fe₇-EDDHA, Gardening Direct, UK) once a week. The working fertilizer solution contained 0.33 g/L Peters Excel 293 Cal-Mag Grower and 0.065 g/L Fe-EDDHA.

### Analysis of gene expression, cloning, construct design and plant transformation

To assess changes in expression of genes encoding the plastid division apparatus, we re-analysed data obtained in a previous study (Hua *et al.,* 2026). Transcript per million were plotted for all genes associated with plastid division and these data used to select specific genes for misexpression. Coding sequences of *ZmG2* (Zm00001d039260), *OsFtsZ1* (LOC_Os04g56970), *OsFtsZ2* (LOC_Os03g44420), *OsPDV1c* (*LOC_Os01g12460*), *OsPDV2* (LOC_Os07g03130), and *OsDRP5B* (LOC_Os12g07880) were synthesized (General Biosystems) and cloned into pDONR223. Coding sequences of *OsMCD1* (LOC_Os04g49540), *OsMINE* (LOC_Os12g31450), and *OsARC6* (LOC_Os02g03000) were amplified from Kitaake cDNA and domesticated for the Golden Gate cloning system. Level-0 modules were then subcloned into level-1 vectors and combined into the level-2 destination vector pICSL4723. Promoters used were the maize *UBIQUITIN* promoter (*ZmUBI_pro_)(Cornejo et al., 1993)*, *SULFITE REDUCTASE* promoter (*OsSiR_pro_)(Hua et al., 2025)*, and the *Zoysia japonica PHOSPHOENOLPYRUVATE CARBOXYKINASE* promoter (*ZjPCK_pro_*)(Nomura *et al*., 2005). Full construct information is provided in Table S1.

Agrobacterium-mediated transformation of Kitaake rice was performed after (Hiei & Komari, 2008) with minor modifications. Dehusked seeds were sterilized with 10% (v/v) bleach for 15 min, rinsed, and cultured on nutrient broth (NB) callus induction medium containing 2 mg/L 2,4-dichlorophenoxyacetic acid at 28 °C in the dark for 3–4 weeks. Actively growing calli were co-cultivated with *A. tumefaciens* strain LBA4404 in NB inoculation medium containing 40 µg/mL acetosyringone for 3 d at 22 °C in the dark. Calli were then transferred to NB recovery medium with 300 mg/L timentin for 1 week at 28 °C in the dark, followed by NB selection medium containing 35 mg/L hygromycin B for 4 weeks at 28 °C in the dark. Resistant calli were moved to NB regeneration medium [100 mg/L myo-inositol, 2 mg/L kinetin, 0.2 mg/L 1-naphthaleneacetic acid (NAA), 0.8 mg/L 6-benzylaminopurine (BAP)] and incubated for 4 weeks at 28 °C in the light. Plantlets were then transferred to NB rooting medium containing 0.1 mg/L NAA for 2 weeks at 28 °C in the light before transfer to soil. Finally, plants were transferred to clay in a walk-in plant growth chamber. Genomic DNA was extracted from T₀ seedlings using a modified CTAB method (Doyle & Doyle, 1987) to confirm transgene presence. Insertion copy number was determined by DNA blotting and TaqMan assays. Two to three independent single-copy T₀ lines were advanced to T₁, and 10–14 plants per line analyzed for gene presence and expression. Homozygous T₂ lines were identified by segregation under 50 mg/L hygromycin. Transgene expression was confirmed with quantitative RT-PCR. Fourth leaves were harvested once fully expanded. RNA was extracted using RNeasy kit (QIAGEN), first-strand cDNA synthesized using SuperScript II reverse transcriptase (Invitrogen) and RT-qPCR performed using SYBR Green PCR Master Mix (Applied Biosystems).

### Scanning electron, confocal laser scanning and brightfield microscopy

Samples were fixed overnight at room temperature in 2% (v/v) glutaraldehyde and 2% (w/v) formaldehyde in 0.05 M sodium cacodylate buffer (pH 7.4) containing 2 mM CaCl₂. Following five washes in 0.05 M sodium cacodylate buffer, samples were osmicated in 1% (w/v) osmium tetroxide and 1.5% (w/v) potassium ferricyanide in 0.05 M sodium cacodylate buffer for 3 d at 4 °C. After rinsing five times in deionized water, samples were treated with 0.1% (w/v) thiocarbohydrazide for 20 min in the dark at room temperature, washed, and re-osmicated in 2% (w/v) osmium tetroxide for 1 h at room temperature. Samples were then washed five times in deionized water and block-stained in 2% (w/v) uranyl acetate (maleate buffer, pH 5.5) for 3 d at 4 °C before washing in deionized water. Dehydration was performed in graded ethanol (50%, 70%, 95%, 100%, 100% (v/v)), followed by 100% acetone and 100% acetonitrile. Samples were infiltrated with 50:50 acetonitrile/Quetol resin overnight, then in 100% Quetol resin for 3 d. A final infiltration was carried out for 5 d in Quetol resin containing BDMA (12 g Quetol 651, 15.7 g NSA, 5.7 g MNA, 0.5 g BDMA; TAAB Laboratories Equipment Ltd), with daily exchanges. Samples were embedded and cured at 60 °C for 2 d. Thin sections (∼100 nm) were then cut using a Leica Ultracut E ultramicrotome, placed on Melinex coverslips and allowed to air-dry. Coverslips were mounted on aluminium SEM stubs using conductive carbon tabs and edges of the slides painted with conductive silver paint. Samples were then sputter-coated with 30 nm carbon (Quorum Q150T E). Imaging was performed on a Verios 460 scanning electron microscope (FEI/Thermofisher) at 4 keV accelerating voltage and 0.2 nA probe current in backscatter mode using the concentric backscatter detector (CBS) in immersion mode at a working distance of 3.5-4 mm; 1536 × 1024-pixel resolution, 3 us dwell time, 4 line integrations. Stitched maps were acquired using FEI MAPS software using the default stitching profile and 5% image overlap. Chloroplast size and number in bundle sheath cells were quantified by confocal imaging (Billakurthi & Hibberd, 2023) with the following modifications. Briefly, the middle region of fourth and seventh leaves were fixed in 1% (v/v) glutaraldehyde (PBS buffer), rinsed twice, and the adaxial surface was gently ablated with a razor blade to expose bundle sheath cells. Tissues were stained with 0.1% calcofluor white (Sigma) for 2 min, rinsed, and imaged on a Leica SP8X upright confocal microscope. Excitation/emission settings: calcofluor white, 405/452–472 nm; chlorophyll autofluorescence, 488/672–692 nm. Images were captured with a 40× water-immersion objective. For each line, three plants were analyzed, with five intermediate veins imaged per plant. Z-stacks were collected from 3–5 bundle sheath cells per vein, yielding data from a minimum of 45 bundle sheath cells per line. Chloroplast planar area, chloroplast number, and bundle sheath cell width, length, and area were quantified using FiJi (Schindelin *et al*., 2012).

To assess chloroplast occupancy with bright field microscopy, cells were isolated after Khoshravesh and Sage (2018). Briefly, leaf tissue was cut into 5-mm × 2-mm strips along the proximodistal axis in a drop of water using a razor blade and immediately immersed in 4% (w/v) paraformaldehyde (pH 6.9; Merck Life Science UK Ltd., Gillingham, UK) at room temperature. Samples were kept in the dark and fixed overnight at 4 °C, after which tissue was transferred to 1× PBS (pH 7.0) and stored at 4 °C. Cell walls were digested by incubating samples in 0.2 M sodium-EDTA (pH 9.0) at 55 °C for 2 h, followed by 2% (w/v) *Aspergillus niger* pectinase (Merck Life Science UK Ltd., Gillingham, UK) at 45 °C for 2 h. Digestion was stopped by incubating tissue in fresh buffer twice for 30 min at room temperature. Individual cells were imaged within 24 h of cell wall digestion. Isolated cells were examined by brightfield microscopy using an Olympus BX51 microscope (Olympus UK and Ireland, Southend-on-Sea, UK). Images were acquired with an MP3.3-RTV-R-CLR-10-C MicroPublisher camera and QCapture Pro 7 software (Teledyne Photometrics, Birmingham, UK). Only bundle sheath cells attached to vascular tissue were selected for imaging. Cell and chloroplast measurements were obtained from scaled images using FiJi (Schindelin *et al*., 2012).

### Chlorophyll quantification, gas exchange and analysis of stomata

The middle region (2 - 3 cm) of fully expanded leaves was harvested, weighed, and immediately flash-frozen in liquid nitrogen. Frozen tissue was ground into a fine powder and suspended in 1 mL of 80% (v/v) acetone. After vortexing, samples were incubated on ice for 15 min with occasional mixing, then centrifuged at 13,000 rpm for 5 min at 4 °C. The supernatant was collected, and the extraction repeated, with pooled supernatants used for analysis. Chlorophyll content was determined spectrophotometrically at 663.6 and 646.6 nm using quartz cuvettes, and concentrations were calculated as described by (Porra *et al*., 1989).

Photosynthetic parameters as assessed by gas exchange were measured on fully expanded 8th or 9th leaves using a LI-6800 portable photosynthesis system (LI-COR Biosciences). Six biological replicates were analysed per line, with each replicate corresponding to an individual plant. Chamber conditions were maintained at 30 °C leaf temperature, 400 µmol s⁻¹ air flow, 60% relative humidity, and 400 µmol mol⁻¹ reference CO₂ concentration. Leaves were acclimated in the chamber for ∼20 min prior to measurements. Net CO₂ assimilation rate (A) in response to intercellular CO₂ concentration (Cᵢ) was determined under a photosynthetic photon flux density (PPFD) of 2,000 µmol photons m⁻² s⁻¹. Reference CO₂ concentrations were sequentially set to 400, 400, 300, 200, 150, 100, 50, 20, 400, 400, 600, 800, 1,000, 1,200, 1,500, and 400 µmol mol⁻¹. Light response curves were generated by measuring A at PPFD values of 2,000, 1,750, 1,500, 1,250, 1,000, 750, 500, 250, 100, 50, 0, and 750 µmol photons m⁻² s⁻¹ at a reference CO₂ concentration of 400 µmol mol⁻¹. The “plantecophys” R package (Duursma, 2015) was used to fit A–Cᵢ curves and estimate the apparent maximum rate of Rubisco carboxylation (Vc_max_) and triose phosphate utilization (TPU). The “photosynthesis” R package was used to determine the light-saturated rate of photosynthesis (k_sat_) and the quantum efficiency of CO₂ assimilation (ϕJ).

Stomatal traits were quantified from the mid-section of fully expanded 8th leaves. Epidermal impressions were obtained by applying a thin layer of clear nail varnish to the abaxial surface, allowing it to dry (∼20 min), and then removing the imprint with transparent tape, which was mounted on a microscope slide. To account for diurnal differences in stomatal aperture, impressions were collected during the light period (10:00–12:00) and the dark period (19:00–21:00), at least one hour after lights were turned off). Images were captured using a Leica SP8X confocal microscope in brightfield mode at 20× magnification. Five images were taken per biological replicate, with 4–6 replicates per genotype. Stomatal length and density were quantified using FiJi (Schindelin *et al*., 2012).

## Results

### Chloroplast development and division gene dynamics during leaf maturation

Consistent with our previous identification of a bundle sheath maturation gradient (Hua *et al*., 2026), light microscopy confirmed that chloroplast size and number increased from the shoot apical meristem toward mature leaf tissues (Fig. 1a-c and Fig. S1a). A significant rise in chloroplast number occurred between stages 3 and 4 in both mesophyll and bundle sheath cells (Fig. 1b). However, the timing of chloroplast expansion was cell-type specific, mesophyll chloroplast size peaked between stages 4 and 5, whereas bundle sheath chloroplast size peaked later, between stages 5 and 6 (Fig. 1c).

**Fig. 1.**
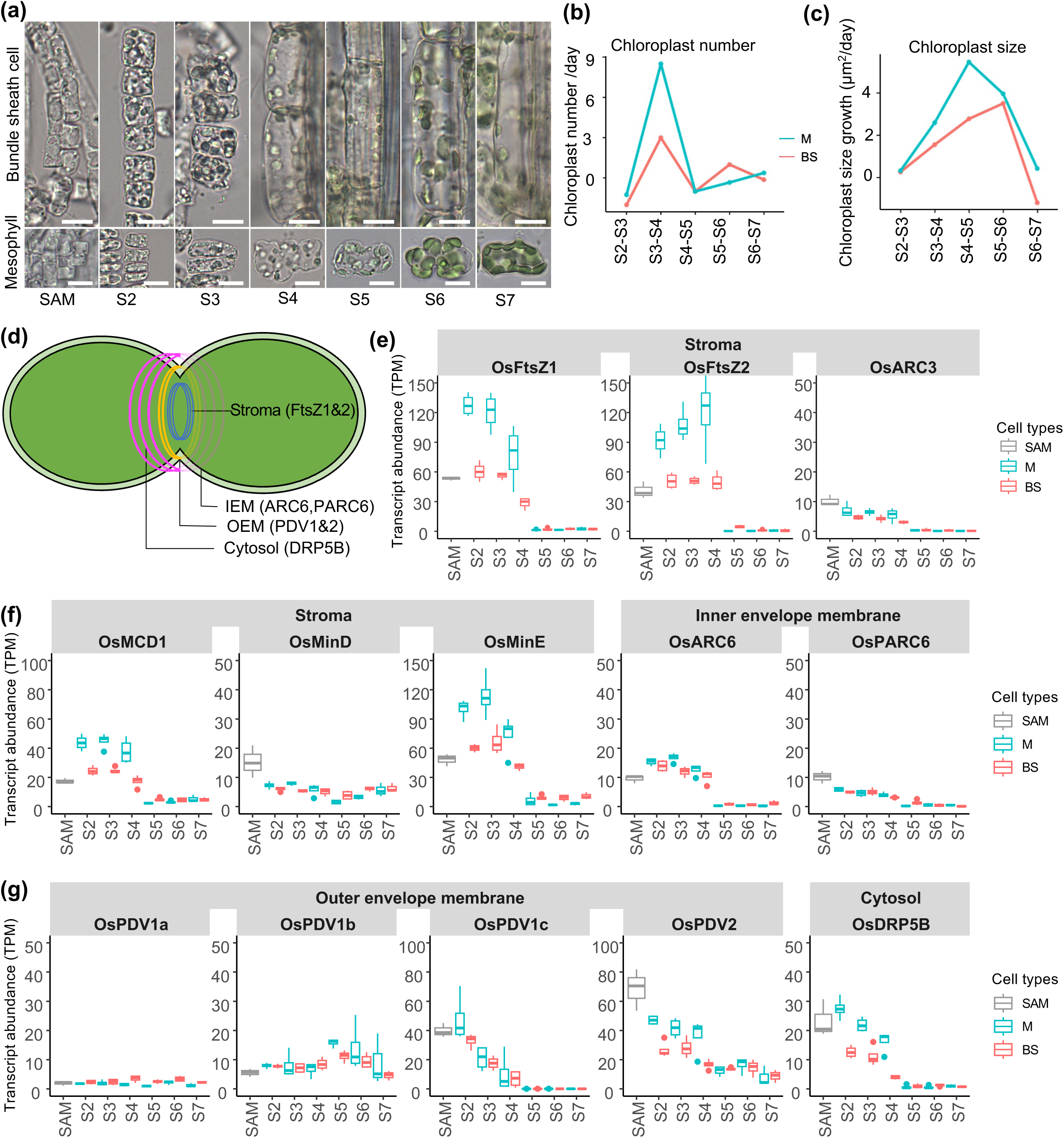
Expression dynamics of chloroplast division genes during development of mesophyll and bundle sheath cells. (a) Chloroplast developmental progression from shoot apical meristem (SAM) to stage 7 (S7) in mesophyll (M) and bundle sheath (BS) cells. (b, c) Rates of increase in chloroplast number (b) and chloroplast size (c) across developmental stages in M and BS cells. (d) Schematic of a dividing chloroplast showing key components of the division machinery. IEM, inner envelope membrane; OEM, outer envelope membrane. (e-g) Expression profiles of chloroplast division genes across developmental stages in M or BS cells. *FtsZ1* and *FtsZ2* assemble the stromal Z-ring; its positioning is regulated by *ARC3*, *MinE*, *MinD* and *MCD1*. *ARC6* and *PARC6* anchor the Z-ring to the IEM, while *PDV1* and *PDV2* recruit the cytosolic dynamin-related protein *DRP5B* at the OEM. *OsFtsZ1*, *OsFtsZ2*, *OsMCD1*, *OsMinE*, *OsARC6*, *OsPDV1c*, *OsPDV2* and *OsDRP5B* showed highest expression during early stages.

Quantification of transcripts encoding core division machinery (*OsFtsZ1, OsFtsZ2, OsMCD1, OsMinE, OsARC6, OsPDV1c, OsPDV2,* and *OsDRP5B* - see Fig 1d) revealed elevated expression during early developmental stages (Fig. 1e-g and Fig. S1b-c). Notably, transcript abundance was generally higher in mesophyll cells than in the bundle sheath at these stages. These data suggest that the cell-specific control of chloroplast proliferation in rice is linked to the differential timing and magnitude of plastid division gene expression.

### Rice chloroplast division genes primarily influence size rather than number

We next tested whether mis-expression of these genes in rice mirrored the phenotypes known in Arabidopsis. For *FtsZ*, as expected, the degree of overexpression varied by line (Fig.S2 and Table S2). In strong overexpression lines, chloroplasts were enlarged and morphologically irregular (Fig. 2a and Fig. S2a-e). While moderate overexpression had no significant effect, strong overexpression reduced bundle sheath chloroplast number by 38% while increasing individual chloroplast area by 50% (Fig. 2b-c). Strong *FtsZ1&2* expression was associated with increased elongation of bundle sheath cells along the proximal-distal leaf axis, but reduced growth along the medial-lateral axis (Fig. S2f–h). Despite these dramatic morphological shifts paralleling dose-dependent phenotypes seen in Arabidopsis, total chlorophyll content remained unaffected (Fig. S2i). For *OsMCD1* and *OsMinE,* because both these genes showed higher natural expression in mesophyll cells during proliferation, we generated co-overexpression lines (Fig. S3 and Table S3). In rice, unlike Arabidopsis, elevated *OsMCD1* and *OsMinE* expression did not have a consistent effect on chloroplast number per cell, individual chloroplast area was reduced in both lines (Fig. 2d-f). Although *PDV* overexpression in Arabidopsis increases plastid number, rice lines overexpressing *OsPDV1c&2* showed no statistically robust change in chloroplast count. Instead, these lines exhibited larger individual chloroplasts and wider bundle sheath cells (Fig. 2g-i, Fig. S4 and Table S4). Lastly, when we overexpressed *OsARC6* and *OsDRP5B,* these constriction ring components also increased chloroplast area by 12–17% without affecting total chloroplast number, or chlorophyll levels (Fig. 2j-l and Fig. S5). In summary, as manipulating the division machinery in rice primarily modulated plastid size rather than number, these combined data demonstrate a significant functional divergence between rice and Arabidopsis.

**Fig. 2.**
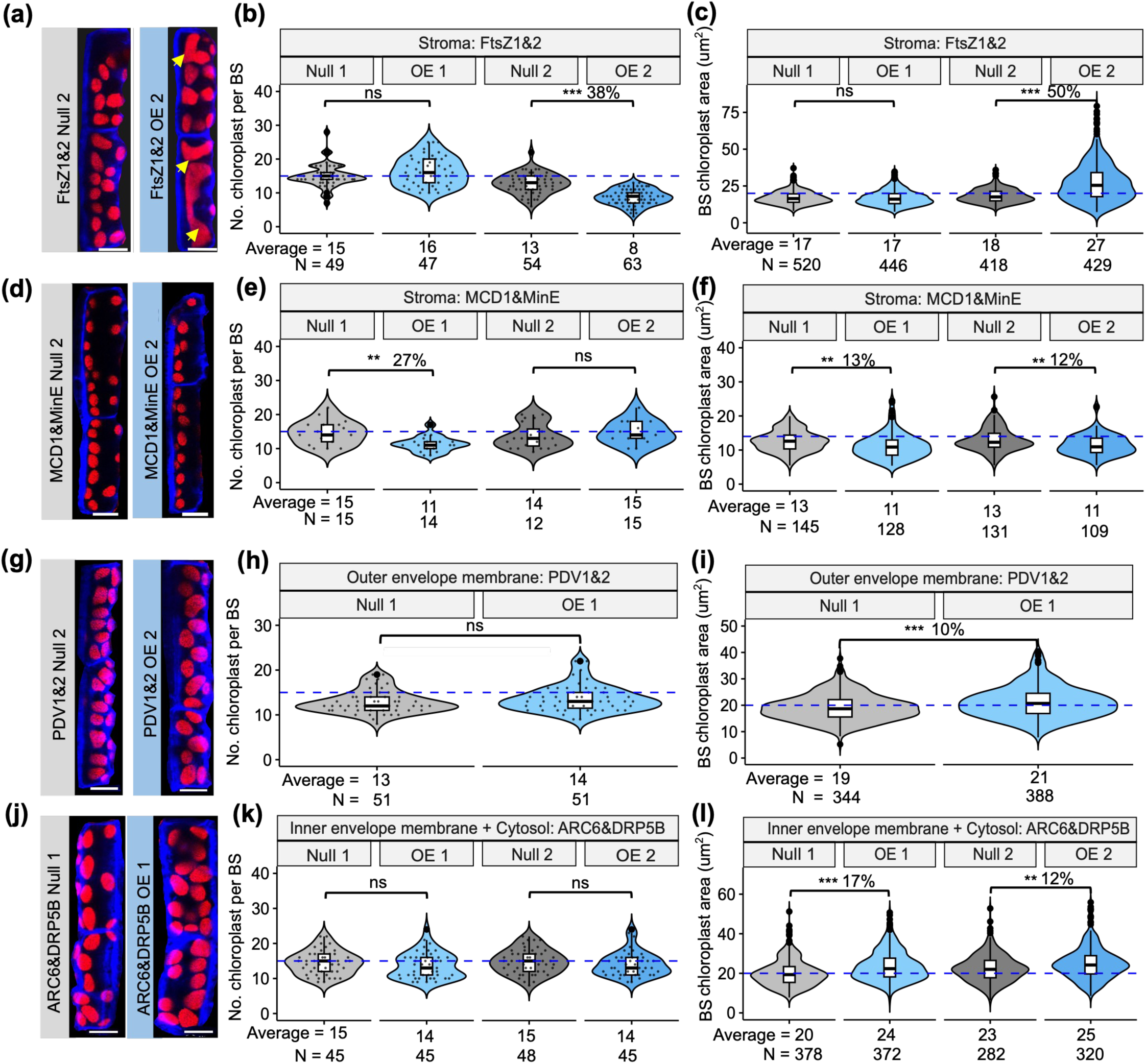
Overexpression of chloroplast division genes alters bundle sheath chloroplast morphology, number, and size in rice. (a-c) *OsFtsZ1* and *OsFtsZ2* overexpression; (a) Confocal images of bundle sheath chloroplasts from the fourth leaf; aberrant and enlarged chloroplasts indicated by yellow arrows. (b) Quantification of chloroplast number per bundle sheath cell in the fourth leaf. (c) Chloroplast area in bundle sheath cells from the fourth leaf (). (d–f) Effects of *OsMCD1* and *OsMinE* overexpression; (d) Confocal images of bundle sheath chloroplasts from the fourth leaf. (e) Chloroplast number per bundle sheath cell. (f) Chloroplast area in bundle sheath cells from the fourth leaf. (g–i) *OsPDV1* and *OsPDV2* overexpression; (g) Confocal images of bundle sheath chloroplasts from the fourth leaf. (h) Chloroplast number per bundle sheath cell. (i) Chloroplast area in bundle sheath cells from the fourth leaf. (j–l) Effects of *OsARC6* and *OsDRP5B* overexpression; (j) Confocal images of bundle sheath chloroplasts from the fourth leaf. (k) Chloroplast number per bundle sheath cell. (l) Chloroplast area in bundle sheath cells from the fourth leaf. Asterisks above violin plots indicate statistically significant differences, as determined by independent *t*-test: *p* ≤ 0.05 (\**), p* ≤ 0.01 (\*\**), p* ≤ 0.001 (***). Non-significant comparisons are labelled as “ns”. Mean chloroplast area values shown in the figure were rounded to the nearest whole number for clarity.

### *ZmG2* and *OsPDV1c&2* co-expression enhances biogenesis but lacks additive effects

We investigated whether combining division genes with the transcription factor *ZmG*2 which is known to promote bundle sheath greening, could further boost chloroplast biogenesis. *ZmG2* combined with *OsARC6&OsDRP5B* did not yield additive effects (Fig. S6). When *ZmG2* was overexpressed with *OsPDV1c* &*OsPDV2* (Fig. S7) larger chloroplasts in the bundle sheath were detected (Fig. 2). Brightfield microscopy confirmed that chloroplasts in bundle sheath cells were larger than in nulls, while mesophyll chloroplast size remained unchanged (Fig. 3a-c). Larger chloroplasts were also visible after confocal laser scanning microscopy (Fig. 3d) and the higher throughput possible with this assay showed that chloroplast area was increased in all lines by 8-23% in both leaf 4 and leaf 7 (Fig. 4d–e; Fig. S8a–b). There was no consistent statistically significant effect on chloroplast number per bundle sheath cell between lines (Fig. 3f). Overall, chloroplast number was comparable between *G2* and the *G2,PDV* lines (Fig.S8c). Scanning electron microscopy revealed normal chloroplast ultrastructure in bundle sheath cells (Fig. 3g; Fig. S9), and also revealed enhanced chloroplast development in the mestome sheath (Fig. 3g). We detected no consistent effect of manipulating chloroplast biogenesis and division on bundle sheath length of width (Fig. 3h-I; Fig. S8d). These findings confirm that *ZmG2* alone and in combination with *OsPDV1c&2* increases chloroplast biogenesis in both the bundle and mestome sheath, but that affects are not additive.

**Fig. 3.**
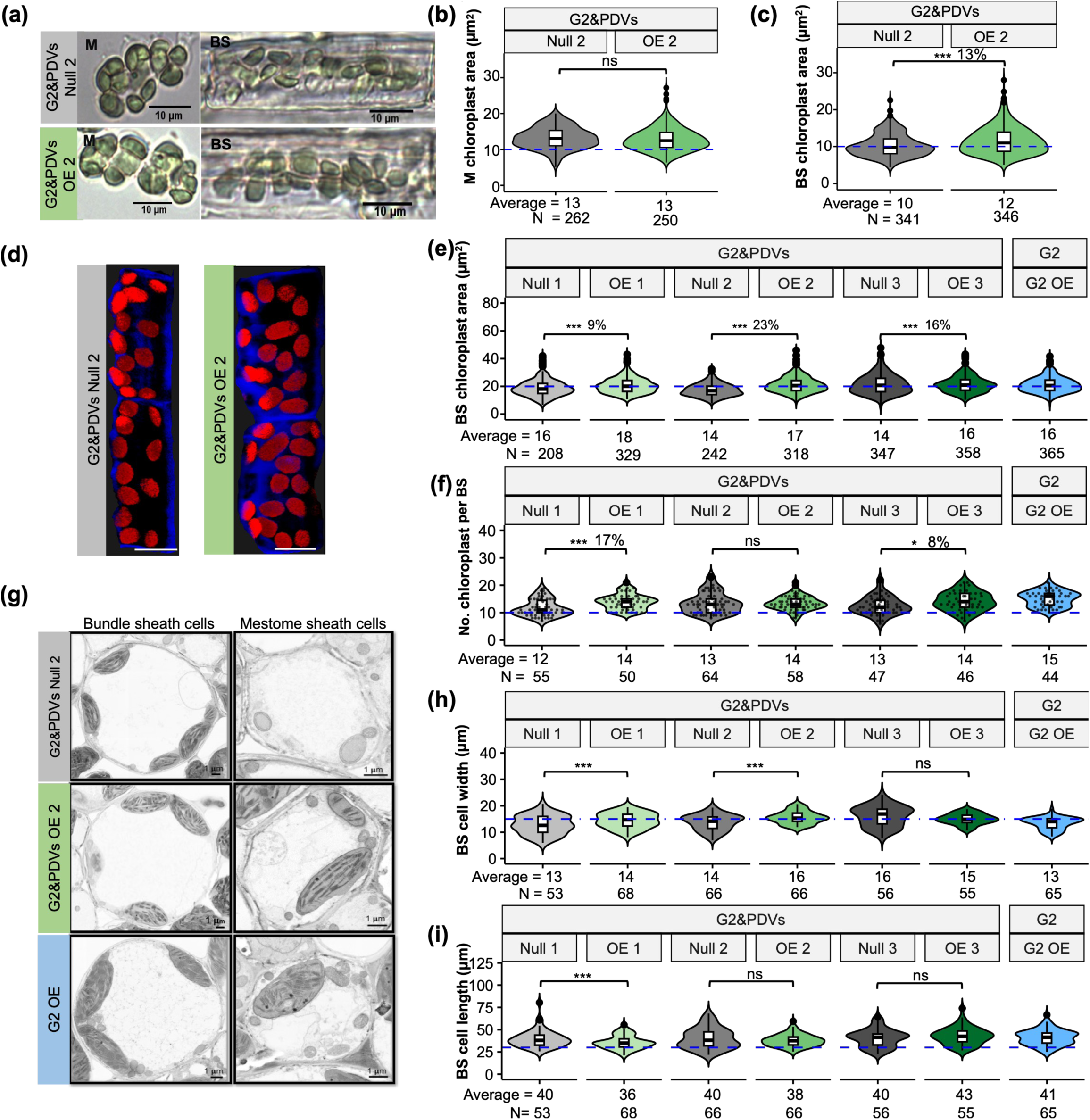
Overexpression of *ZmG2*, *OsPDV1*, and *OsPDV2* increases chloroplast size in bundle sheath cells. (a) Representative brightfield microscopy images of mesophyll (M) and bundle sheath (BS) cells. (b, c) Quantification of chloroplast area in mesophyll and bundle sheath cells from brightfield microscopy. (d) Representative confocal images of bundle sheath cell chloroplasts. (e) Quantification of chloroplast area in bundle sheath cells of leaf 7 from confocal images. (f) Quantification of chloroplast number per bundle sheath cell from confocal images. (g) Scanning electron microscopy images of chloroplasts in bundle sheath and mesophyll cells. (h, i) Bundle sheath cell width and length in leaf 4, quantified from confocal images. Stars above violin plots indicate statistically significant differences between overexpression lines and corresponding nulls, as determined by independent *t*-test: *p* ≤ 0.05 (\**), p* ≤ 0.01 (\*\**), p* ≤ 0.001 (***). Non-significant comparisons are labelled as “ns”. Mean chloroplast area values shown in the figure were rounded to the nearest whole number for clarity.

**Fig. 4.**
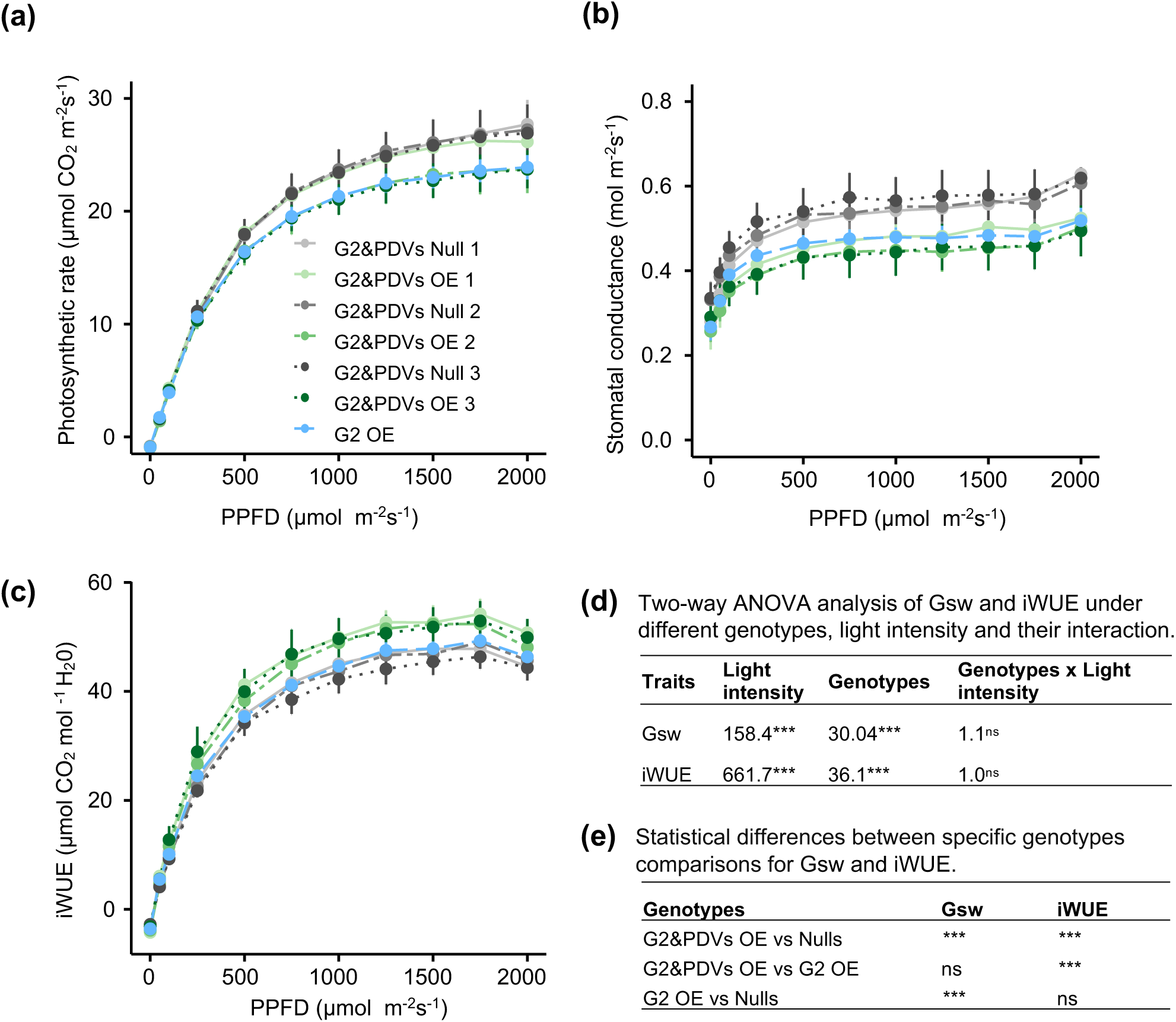
Overexpression of *ZmG2*, *OsPDV1*, and *OsPDV2* did not affect photosynthesis rate but enhanced water use efficiency in rice. (a) Light response curves for each genotype – statistical testing indicated no significant differences. (b) Stomatal conductance (Gsw) was statistically reduced in overexpression lines (see d&e). (c) Intrinsic water use efficiency (iWUE) was increased in overexpression lines. (d) Two-way ANOVA analysis of Gsw and iWUE for each genotype, light intensity and their interaction. (e) Statistical differences between specific genotypes comparisons for Gsw and iWUE as determined by independent *t*-test: *p* ≤ 0.05 (\**), p* ≤ 0.01 (\*\**), p* ≤ 0.001 (***). Non-significant comparisons are labelled as “ns”.

### Combined overexpression improves water-use efficiency via stomatal remodelling

Finally, we assessed whether the changes in bundle sheath chloroplast volume documented above for the *FtzS* and *PDV* lines impacted leaf-level physiology. In *FtsZ1&2* lines, photosynthetic rates and stomatal conductance were largely unaffected (Fig. S10a-e). Photosynthetic performance however did decline under at elevated CO₂ in the strong overexpression line (Fig. S10f-h). The *ZmG2,OsPDV1&2* lines however did exhibit distinct profiles (Fig. 4 and Fig.S11). For example, although light-response and A/Ci analyses showed no statistically robust differences between genotypes (Fig. 4a; Fig. S11d–i), we did detect significant effects of both light intensity and genotype on stomatal conductance and intrinsic water-use efficiency (two-way ANOVA; gsw: F = 158.4, *p* < 0.001; iWUE: F = 661.7, *p* < 0.001; Fig. 4b-e). Specifically, overexpression of both *G2* and *PDVs* led to lower stomatal conductance than nulls and produced higher instantaneous water use efficiency compared with both *G2*-only and nulls (Fig.4b-e). Under both growth and saturating light higher instantaneous water use efficiency was sustained (Fig. S12; Tables S5–S6). This increase in instantaneous water use efficiency was associated with anatomical changes in the epidermis. In the Zm*G2, PDV* lines, stomatal density increased, and individual stomata were significantly shorter (Fig. 5a-d and Fig.S13). These results suggest that manipulating chloroplast division and biogenesis factors in rice may trigger unexpected developmental crosstalk that impacts stomatal structure and water-use efficiency.

**Fig. 5.**
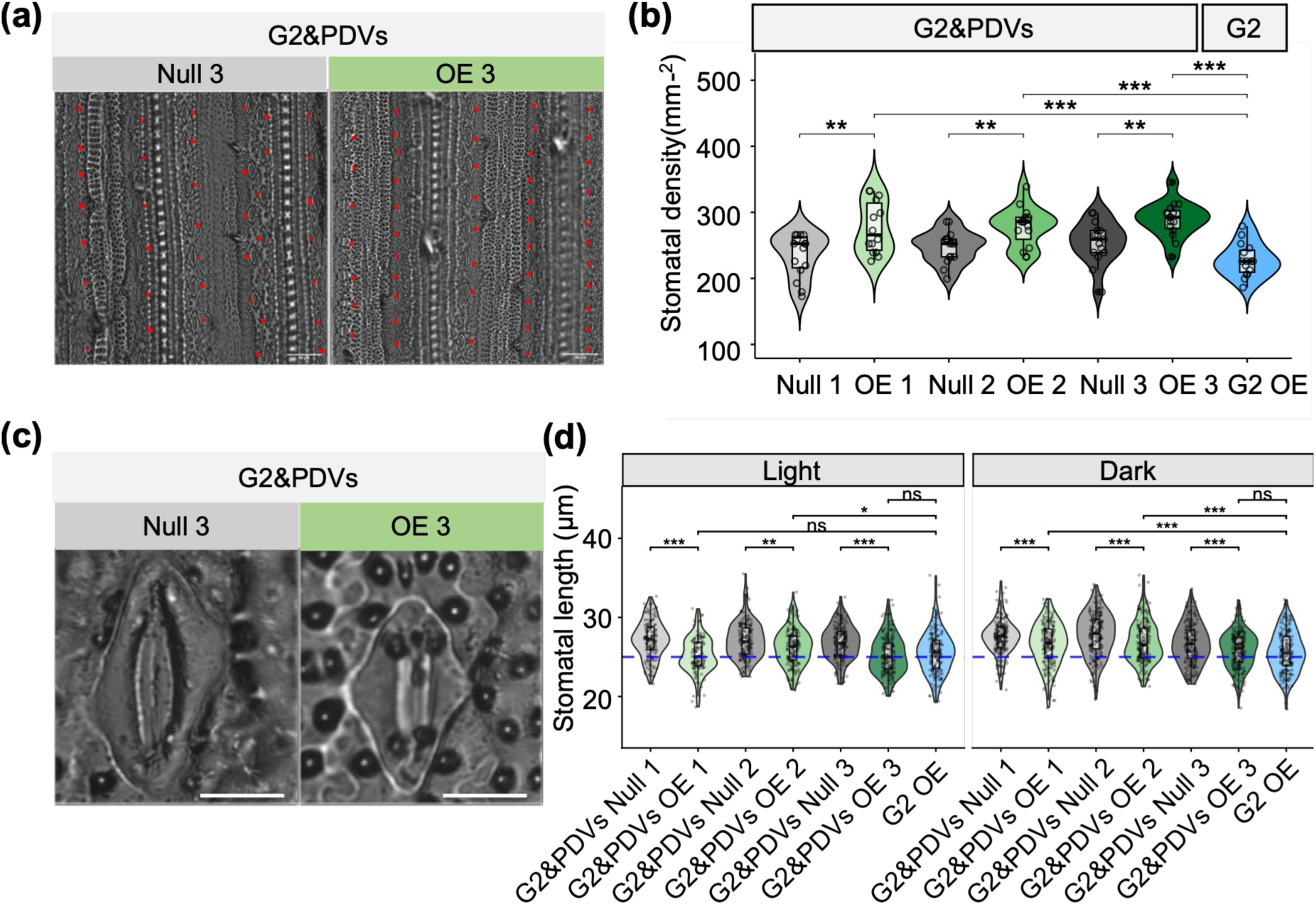
Overexpression of *ZmG2*, *OsPDV1*, and *OsPDV2* increases stomatal density and reduces stomatal length. (a) Images of leaf surface with stomatal indicated by red dots (bar scale = 50 µm). (b) Quantification of stomatal density, showing statistically significant differences between overexpression lines *of ZmG2, OsPDV1,* and *OsPDV2* compared with controls. (c) Images of stomata from the adaxial side (bar sale=10 µm). (d) Stomatal length under light and dark conditions on the abaxial leaf blade showing reductions in overexpression lines. Stars above violin plots indicate statistically significant differences between lines, as determined by independent *t*-test: *p* ≤ 0.05 (\**), p* ≤ 0.01 (\*\**), p* ≤ 0.001 (***). Non-significant comparisons are labelled as “ns”.

## Discussion

### A precise developmental window for plastid engineering and functional divergence of the rice division machinery

Our analysis defines a discrete developmental window between stages 3 and 4 of leaf expansion as defined by our sampling where chloroplast proliferation is most active and core division gene expression peaks (Fig. 1). Interestingly, we observed a temporal offset in maturation, with mesophyll plastids expanding earlier than those in the bundle sheath. Similar offsets have been reported in C_4_ species such as maize and *Setaria* (Majeran *et al*., 2005; Majeran & van Wijk, 2009; John *et al*., 2014). This conserved heterochrony suggests that hallmarks of the developmental “blueprint” for dual-cell-type plastid maturation are present in the C_3_ model rice. For biotechnological applications, our findings emphasize that interventions to increase plastid number likely need to be synchronized with this early proliferation phase to be effective.

A central finding of this study is that the phenotypic consequences of manipulating chloroplast division genes in rice differ significantly from those established in *Arabidopsis*. While strong *OsFtsZ1/2* overexpression produced the classic “giant chloroplast” phenotype, other components showed distinct behaviours. For example, when *MinE* that determines ring positioning was overexpressed in *Arabidopsis* division was inhibited and plastids were larger. In rice, however, *OsMinE&OsMCD1* overexpression restricted expansion, resulting in smaller plastids. Notably, overexpressing *OsPDV1&2* or *OsARC6&DRP5B* in rice increased plastid size rather than number. This directly contrasts with *Arabidopsis*, where *PDV* overexpression stimulates division to produce more, smaller chloroplasts (Okazaki *et al*., 2009). These discrepancies suggest that the stoichiometric requirements for the division machinery are more stringent in monocotyledons, or that downstream recruitment of the constriction ring is limited by different bottlenecks. Our results underscore that simply “tuning up” individual structural components is insufficient to drive proliferation; rather, it perturbs the delicate balance of the division site, leading to morphological defects to the chloroplast rather than increased plastid counts.

### Coordinated improvement of chloroplast occupancy and water-use efficiency

An unexpected but interesting outcome of this study was the physiological shift observed in lines in which chloroplast biogenesis was increased in the bundle sheath after misexpression of both *ZmG2* and *PDV* genes. Previous work showed that constitutive overexpression of *ZmG2* increases stomatal density, reduces guard cell length, and activates ABA-related stress pathways to improve drought tolerance in rice (Li *et al*., 2024). When we also expressed *ZmG2* from the constitutive *UBQ* promoter, but drove *PDVs* from the *ZjPCK* promoter, that is active in both bundle sheath and guard cells (Danila *et al*., 2022), we also found a “stomatal remodelling” phenotype. Specifically, these plants exhibited higher stomatal density, but smaller guard cells compared with controls, a configuration that reduced stomatal conductance without penalising CO_2_ assimilation. This coordinated shift led to a significant increase in intrinsic water-use efficiency. The expression of both *ZmG2* and the *PDVs* had the greatest impact on these traits suggesting a potential link between the plastid division apparatus and stomatal development. Future work could investigate whether this type of manipulation is advantageous in field conditions, and whether there is crosstalk between PDV proteins and guard cell mother cell divisions, or a retrograde signal from altered bundle sheath plastids.

### Implications for C_4_ engineering and climate resilience

Achieving C_4_-like chloroplast occupancy in rice is a multifaceted challenge. Our results demonstrate that while structural division factors (like FtsZ and PDVs) can modulate plastid volume, they do not independently drive the massive increase in biogenesis required for a functional C_4_ cycle. In contrast, the transcription factor *ZmG2* provides a superior foundation by promoting differentiation across both the bundle and mestome sheaths (Wang *et al*., 2017). Modifying stomatal traits to improve water-use efficiency without reducing carbon gain is a primary goal for climate-resilient agriculture (Caine *et al*., 2019). The additive benefit of combining *ZmG2* and *PDV* misexpression in terms of improvement in instantaneous water use efficiency presents an avenue that could be explored for crop improvement.

## Acknowledgements

This work was funded by a C_4_ Rice project grant from the Bill Gates Foundation to University of Oxford (INV-002970). We acknowledge Dr. Karin Mueller from the Cambridge Advanced Imaging centre for her help with SEM imaging.

## Acknowledgements

None declared.

## Authors Contributions

CC designed and conducted experiments, analysed data and drafted the manuscript. LH designed, conducted and analysed laser capture microdissection for mesophyll and bundle sheath cells. ARGP developed the staged sampling used to sample development of mesophyll and bundle sheath cells. ARB carried out T0 and T1 plant propagation and phenotyping for overexpression lines of *ZmG2, OsPDV1&V2*. NW, RD and SS conducted rice transformation, and NW undertook the DNA blot analysis. KB designed constructs for *OsFtsZ1&2*. TSc and ARB performed SEM experiments. TSu assisted with plant growth and maintenance. CC and JMH wrote the manuscript with input from all authors. JMH supervised the execution of experiments and oversaw the project.

