## Supplementary material for "Temporal dynamics and functional divergence of the chloroplast division apparatus in *Oryza sativa*": S Figures

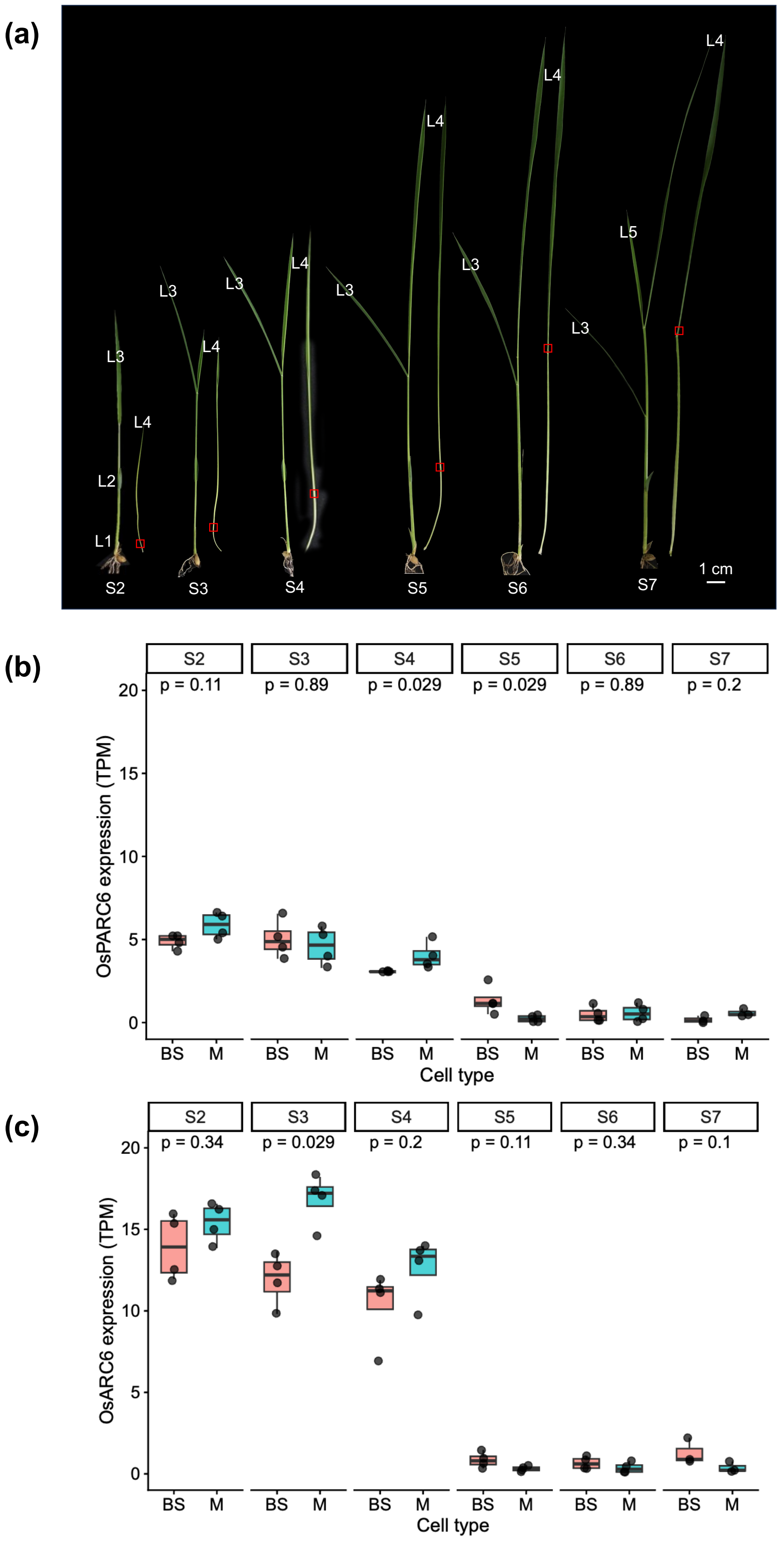


**Fig. S1 Developmental stages of rice leaves from stage 2 to stage 7.** (**a**) Red highlighted areas indicate regions used for quantification of chloroplast size in mesophyll and bundle sheath cells, as well as for laser capture microdissection (LCM) followed by transcriptome sequencing. (**b-c**), Expression profiles of chloroplast division genes, *OsARC6* (b) and *OsPARC6* (b) across developmental stages in M and BS cells. Statistical significance between mesophyll and bundle sheath cells, determined by the Wilcox Test.

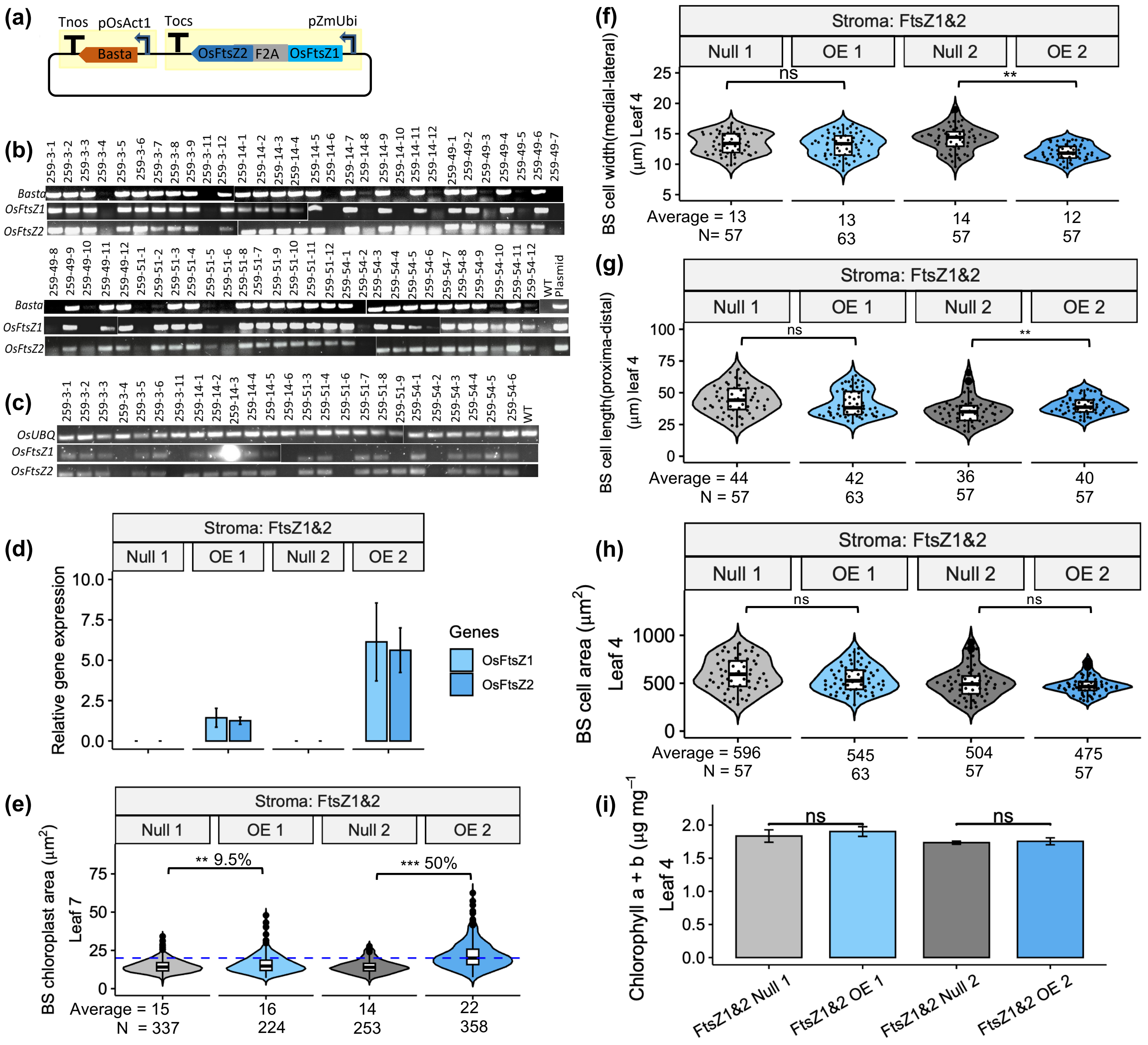

**Fig. S2 Overexpression of *OsFtsZ1* and *OsFtsZ2* at high levels induces abnormal chloroplast morphology and a reduced chloroplast number in rice bundle sheath cells, resulting in a lower photosynthetic rate under high light conditions. (a)** Schematic of the transformation construct. The maize *UBIQUITIN* promoter (*ZmUBI*) was used to drive co-expression of *OsFtsZ1* and *OsFtsZ2*. **(b)** Genotyping of T1 plants from six independent transformation events (259-3, 259-14, 259-49, 259-51, and 259-54), each confirmed to carry a single-copy transgene insertion by by TaqMan assays in the T0 generation. Line 259-14-6 represents the *FtsZ1&2* Null 1 and 259-14-7 is the corresponding overexpression line *FtsZ1&2* OE 1). Likewise, 259-51-6 is *FtsZ1&2* Null 2, and 259-51-8 is *FtsZ1&2* OE 2. **(c)** RT-PCR validation confirms expression of *OsFtsZ1* and *OsFtsZ2* in T1 overexpression lines, with no detectable expression in the respective null segregants. **(d)** qRT-PCR analysis of *OsFtsZ1* and *OsFtsZ2* transcript levels in T2 homozygous overexpression lines and corresponding sibling nulls. **(e)** Quantification of chloroplast area in bundle sheath cells from the fourth leaves. **(f-h):** Quantification of bundle sheath cell length (f), width (g) and size (h). **(i)** Chlorophyll content of leaf 4. Stars above violin plots indicate statistical significance between overexpression lines and corresponding nulls, determined by independent *t*-test: $p \leq 0.05$ (*), $p \leq 0.01$ (**), $p \leq 0.001$ (***); "ns" indicates no significant difference.

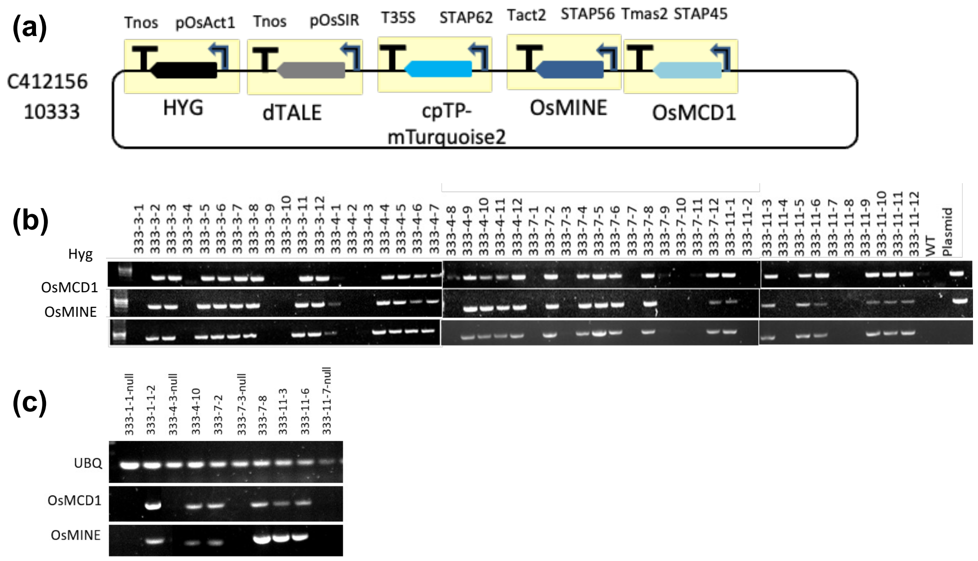


**Fig.S3 Validation of transgenic rice lines after overexpressing *OsMCD1* and *OsMINE*. (a)** Schematic diagram of the transformation construct. The bundle sheath-specific promoter *OsSiR* was used to drive expression of *OsMINE* and *OsMCD1* via the dTALE system. **(b)** Genotyping results of T1 plants from four independent lines (3, 4, 7, and 11), with each confirmed to carry a single-copy transgene insertion by TaqMan assays in T0 generation plants. **(c)** RT-PCR analysis of T1 homozygous overexpression (OE) and corresponding null lines. Line 333-7-3-null corresponds to *MCD1&MINE* Null 1, and 333-7-2-homo corresponds to *MCD1&MINE* OE 1. Line 333-11-7-null corresponds to Null 2, and 333-11-6-homo corresponds to OE 2.

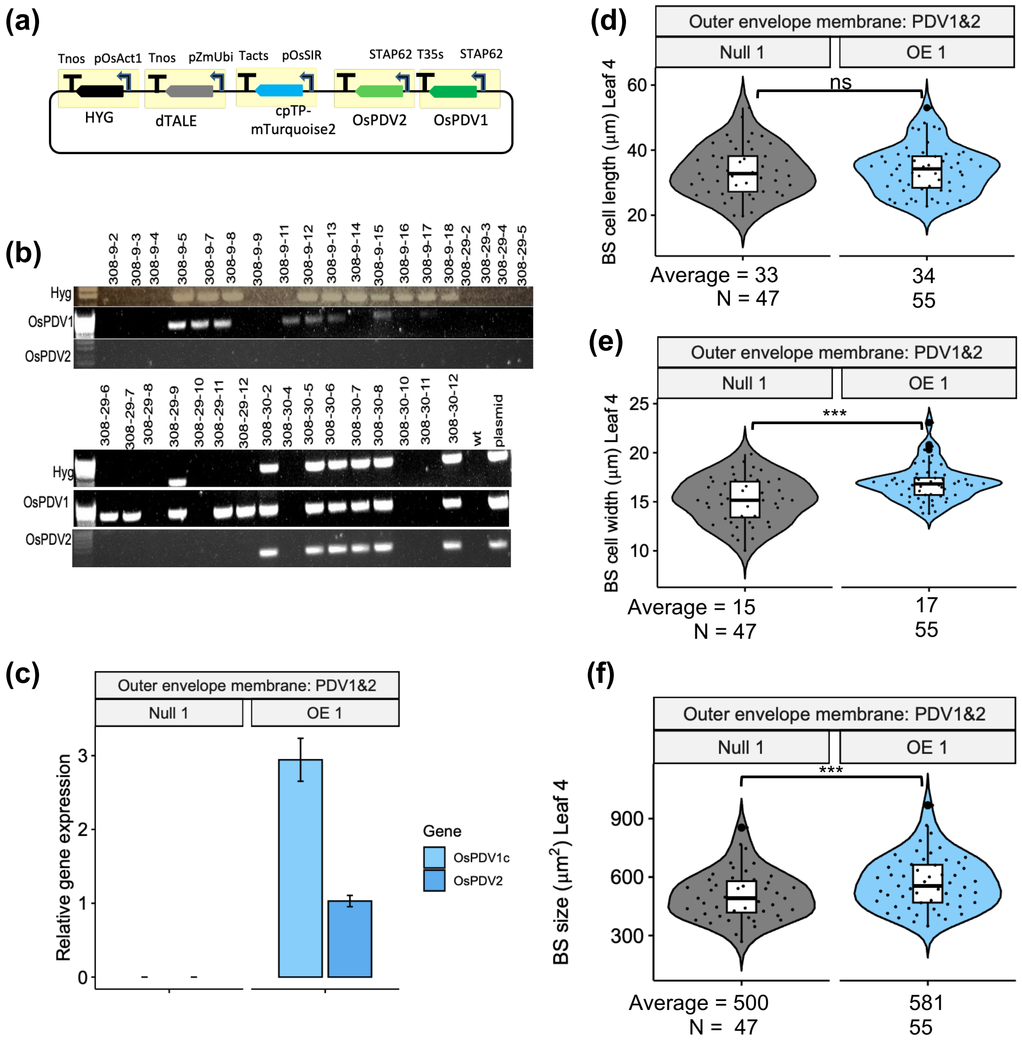


**Fig. S4 Overexpression of *OsPDV1* and *OsPDV2* increases chloroplast and bundle sheath cell size without altering chloroplast number. (a)** Schematic of the transformation construct. The maize *UBIQUITIN* promoter (ZmUbi) was used to drive co-expression of *OsPDV1* and *OsPDV2*. **(b)** Genotyping of T1 plants from three independent transformation events (308-9, 308-29, 308-30), each confirmed to carry a single-copy transgene insertion by digital PCR in the T0 generation. Line 308-30-8 is *PDV1&2* OE 1, and 308-30-10 corresponds to *PDV1&2* Null 1. **(c)** qRT-PCR analysis of *OsPDV1* and *OsPDV2* transcript levels in the T2 homozygous overexpression line and corresponding sibling null. Three biological replicates were used for each line. **(d-f)** Quantification of bundle sheath cell length (d), width (e), and total cell area (f). Stars above violin plots indicate statistical significance between overexpression lines and corresponding nulls, determined by independent *t*-test: $p \leq 0.05$ (*), $p \leq 0.01$ (**), $p \leq 0.001$ (***); "ns" indicates no significant difference.

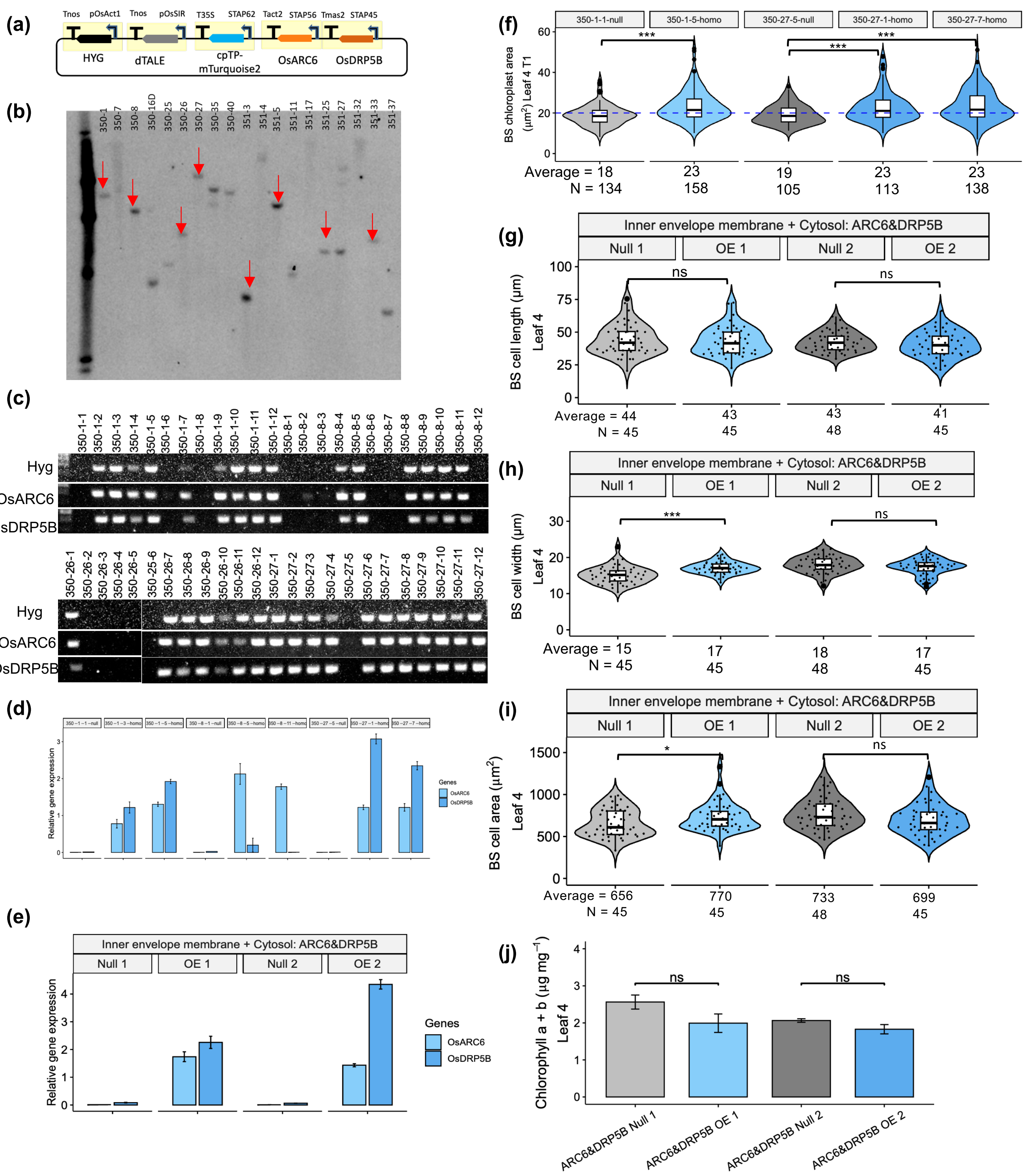


**Fig. S5 Overexpression of *OsARC6* and *OsDRP5B* increase chloroplast size but did not change bundle sheath cell and chlorophyll content.** **(a)** Schematic of the transformation construct. The bundle sheath-specific promoter *OsSiR* was used to drive expression of *OsARC6* and *OsDRP5B* via the dTALE system. **(b)** Southern blot analysis of genomic DNA from $T_0$ transformants identified lines carrying single-copy T-DNA insertions. Four lines from construct 350 (350-1, 350-8, 350-26, and 350-27) and four lines from construct 351 (351-3, 351-5, 351-25, and 351-33) were advanced to the $T_1$ generation. **(c)** Genotyping of $T_1$ plants derived from selected transformation events of constructs 350. Line 350-1-1-null represents the *ARC6&OsDRP5B* null (Null 1), and 350-1-3-homo is the corresponding overexpression line (OE 1). Likewise, 350-25-5-null is Null 2, and 350-27-7 is OE 2. **(d)** Quantitative RT-PCR analysis showing expression of *OsARC6* and *OsDRP5B* in $T_1$ overexpression lines, with no detectable expression in null segregants. **(e)** qRT-PCR analysis of *OsARC6* and *OsDRP5B* transcript levels in the T2 homozygous overexpression line and corresponding sibling null. **(f)** Quantification of chloroplast area in bundle sheath cells from the fourth leaf in T1 plants. 350-1-1-null represents the *ARC6&OsDRP5B* null (Null 1), 350-1-5-homo is the corresponding overexpression line, Likewise, 350-25-5-null is Null 2, and 350-27-1 and 350-27-7 are the corresponding overexpression homozygous lines. **(g-i)** Quantification of bundle sheath cell width (g), length (h), and cell area (i) from T2 homozygous lines of leaf 4. Three biological replicates were used for each line. **(j)** Chlorophyll content of leaf 4. Stars above violin plots indicate statistical significance between overexpression lines and corresponding nulls, determined by independent *t*-test: $p \leq 0.05$ (*), $p \leq 0.01$ (**), $p \leq 0.001$ (***); "ns" indicates no significant difference.

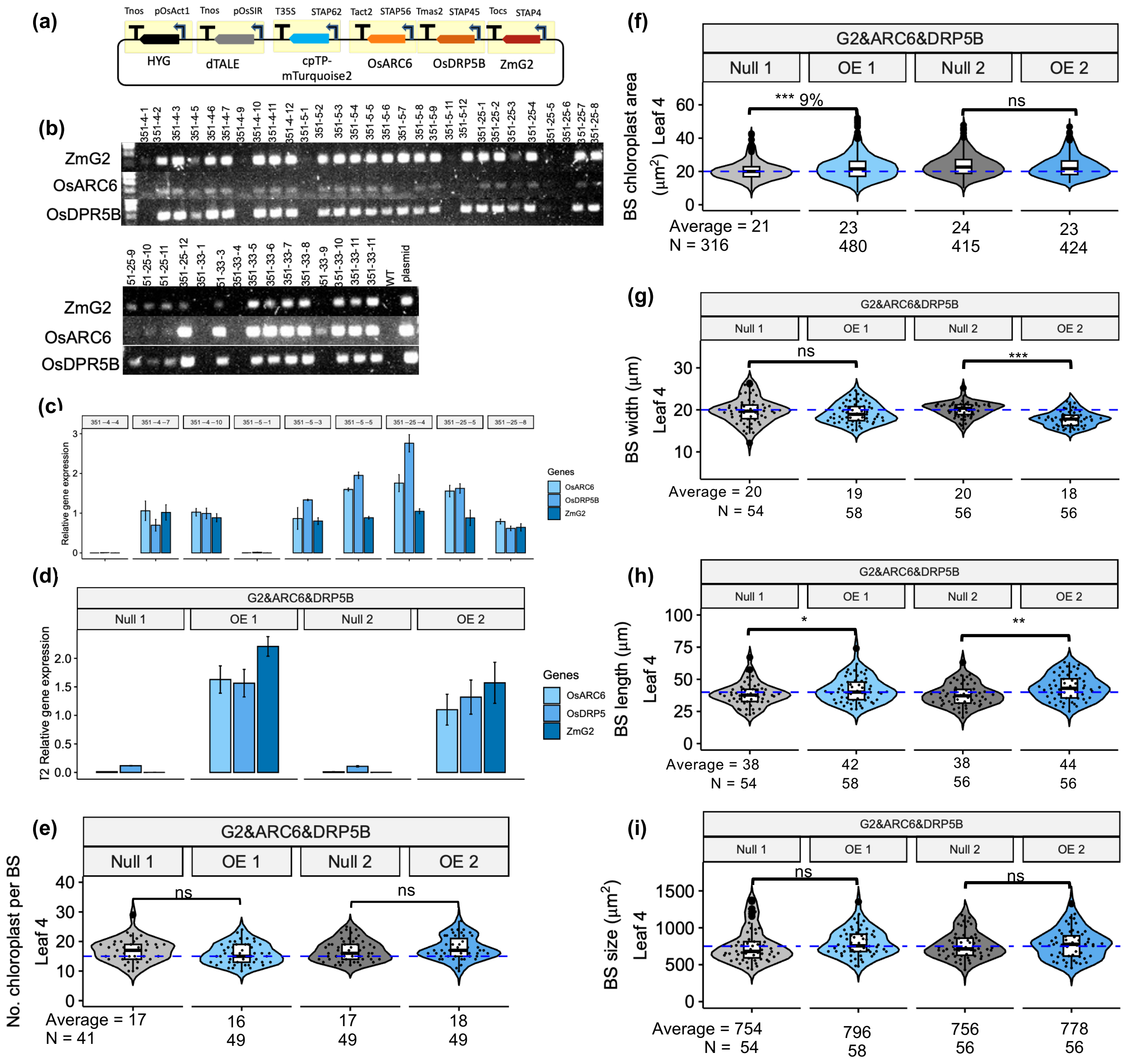


**Fig. S6 Validation of transgenic rice lines overexpressing of *ZmG2, OsARC6* and *OsDRP5B.*** **(a)** Schematic of the transformation construct. The bundle sheath-specific promoter *OsSiR* was used to drive expression of *ZmG2, OsARC6* and *OsDRP5B* via the dTALE system. **(b)** Genotyping of $T_1$ plants derived from selected transformation events of construct 351. Line 351-4-4-null represents the *ZmG2&ARC6&OsDRP5B* null (Null 1), and 351-4-10-homo is the corresponding overexpression line (OE 1). Likewise, 351-5-1is Null 2, and 351-5-5 is OE 2, 351-25-6 is Null 3 and 351-25-8 is OE 3. **(c)** Quantitative RT-PCR analysis showing expression of *ZmG2*, *OsARC6* and *OsDRP5B* in $T_1$ overexpression lines, with no detectable expression in null segregants. **(d)** qRT-PCR analysis of *ZmG2, OsARC6* and *OsDRP5B* transcript levels in the T2 homozygous overexpression line and corresponding sibling null. **(e, f)** Quantification of chloroplast number (a) and chloroplast area (b) in bundle sheath cells from homozygous lines of leaf 4. **(g-i)** Quantification of bundle sheath cell width(g), length (h), and cell area (i) from homozygous lines of leaf 4. Stars above violin plots indicate statistical significance between overexpression lines and corresponding nulls, determined by independent *t*-test: $p \leq 0.05$ (*), $p \leq 0.01$ (**), $p \leq 0.001$ (**); "ns" indicates no significant difference.

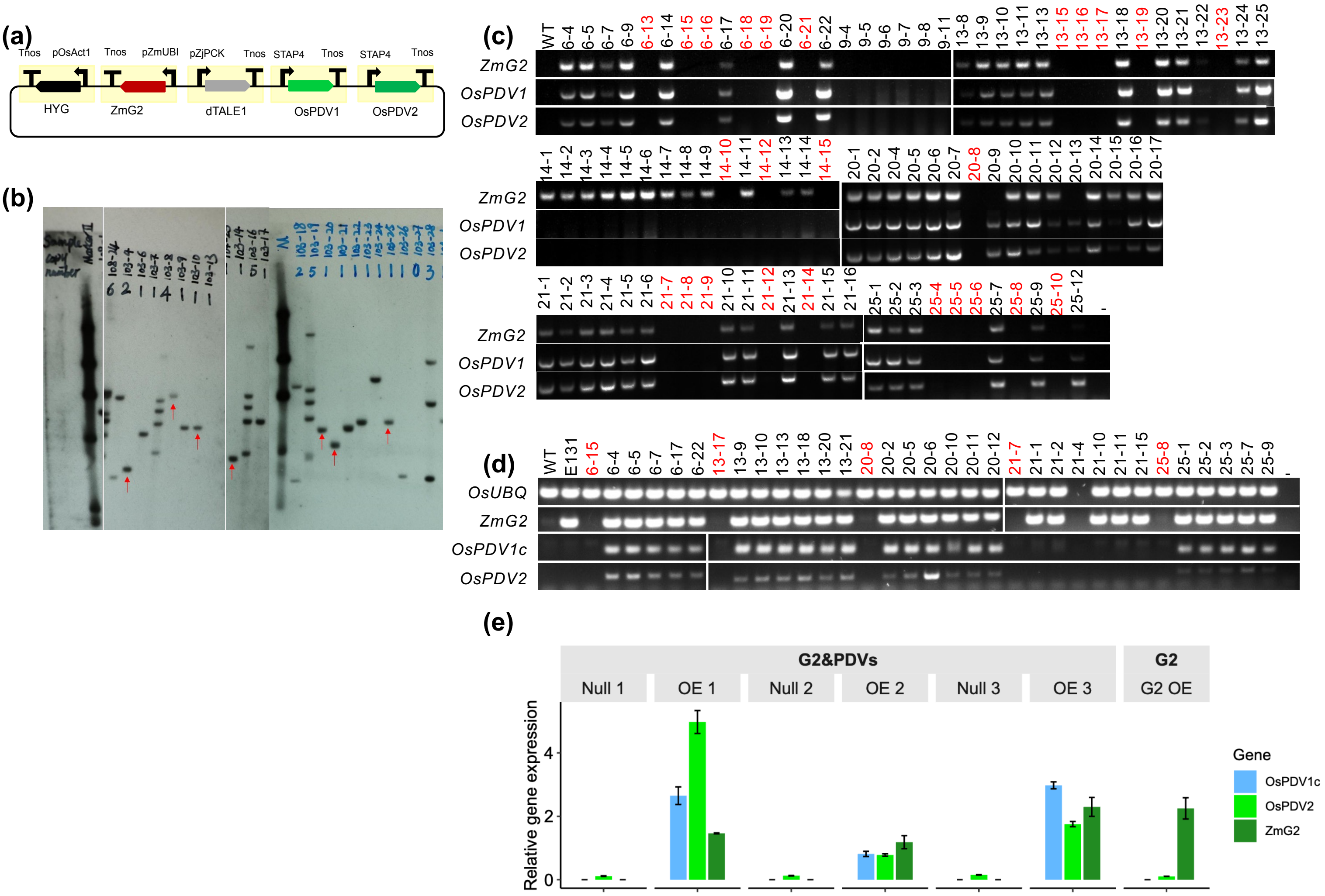


**Fig. S7 Generation and validation of transgenic rice lines overexpressing *ZmG2*, *OsPDV1*, and *OsPDV2*. (a)** Schematic of transformation construct. The maize *UBIQUITIN* promoter (*UBIpro*) was used to drive *ZmG2*, and the *PHOSPHOENOLPYRUVATE CARBOXYKINASE* promoter from *Zoysia japonica* (*ZjPCKpro*), which confers bundle sheath cell–preferential expression, was used to drive *OsPDV1* and *OsPDV2*. **(b)** Southern blot analysis of genomic DNA from $T_0$ transformants was performed to identify lines with single-copy T-DNA insertions. Seven lines (marked with red arrows) carrying single insertions were selected for progression to the $T_1$ generation. **(c)** Genotyping of $T_1$ plants derived from seven independent transformation events (lines 6, 9, 13, 14, 20, 21, and 25) was conducted to identify null segregants (highlighted in red). **(d)** RT-PCR validation of $T_1$ individuals confirmed transgene expression. Line 6-15 represents the *G2&PDVs* Null 1, and 6-5 is the corresponding overexpression line *G2&PDVs* OE 1, Likewise, 20-8 is Null 2, and 20-5 is OE 2, 25-8 is the Null 3 and 25-7 is the OE 3. **(e)** qRT-PCR quantification of *ZmG2*, *OsPDV1*, and *OsPDV2* transcript levels in homozygous overexpression lines and their corresponding sibling nulls.

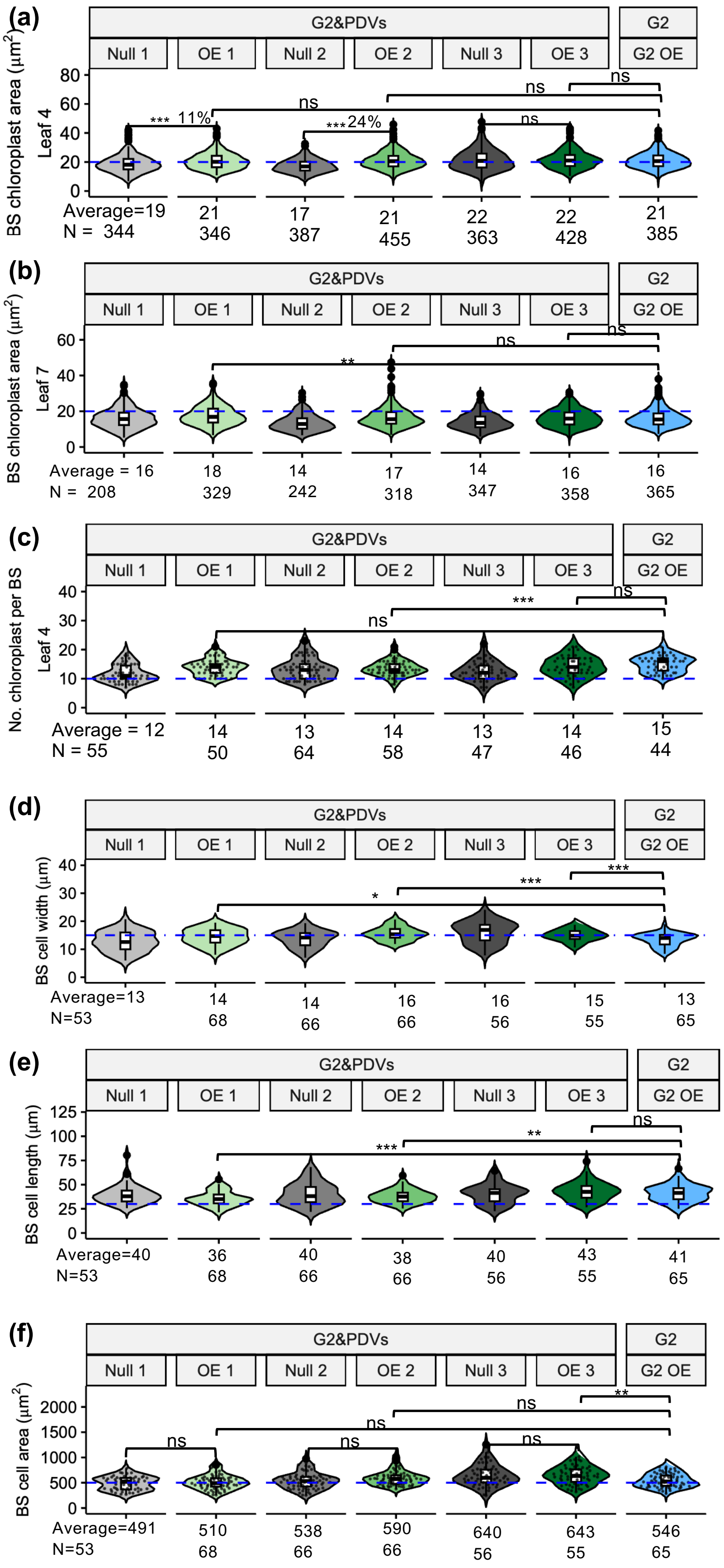


**Fig. S8 Overexpression of *ZmG2*, *OsPDV1*, and *OsPDV2* increases chloroplast size in bundle sheath cells without altering bundle sheath cell size. (a-c)** Quantification of chloroplast area in leaf 4 (a) and leaf 7 (b) and chloroplast number in leaf 4 (c) in bundle sheath cells based on confocal microscopy. **(d-f)** Measurement of BS cell width(d), length (e) and size (f) quantified from confocal images. Stars above violin plots indicate statistical significance between overexpression lines and corresponding nulls, determined by independent *t*-test: $p \leq 0.05$ (*), $p \leq 0.01$ (**), $p \leq 0.001$ (***); "ns" indicates no significant difference.

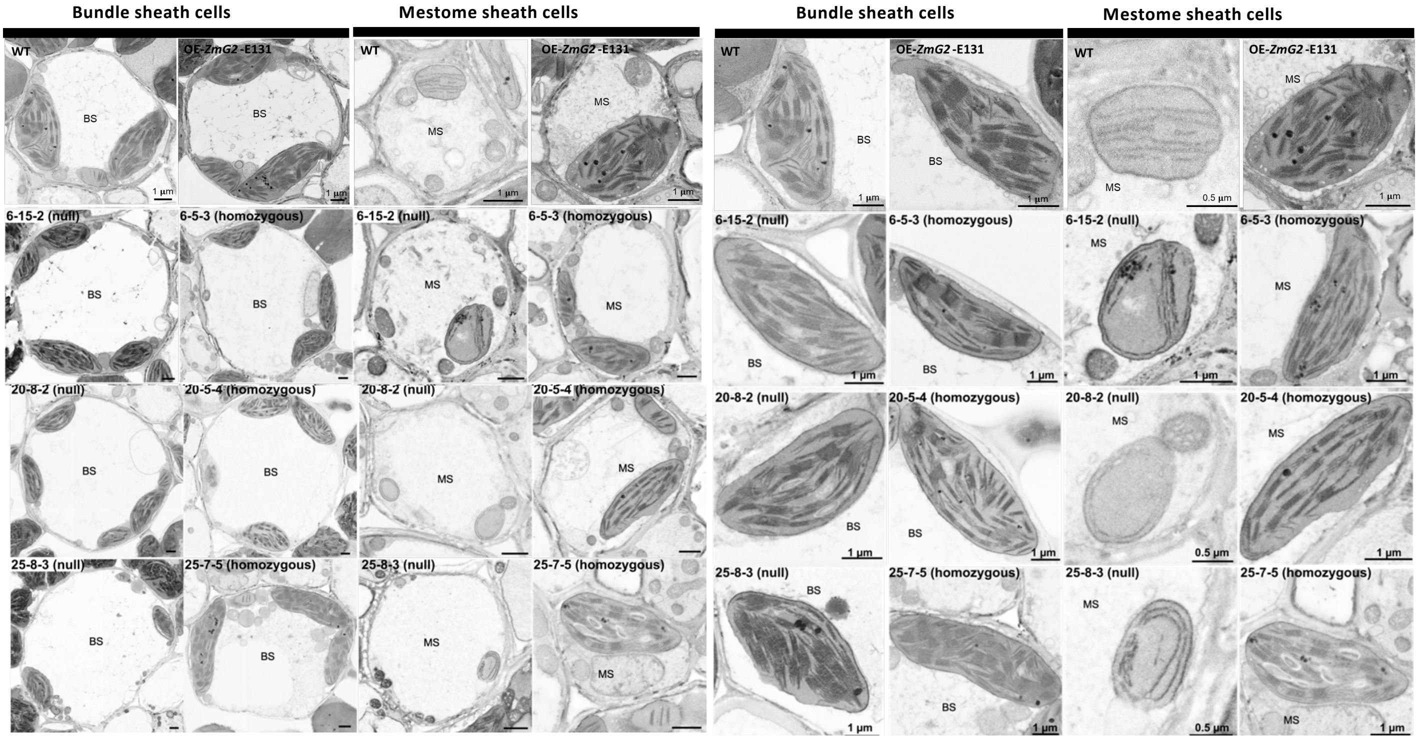


**Fig. S9 Chloroplast ultrastructure by SEM from minor veins of leaf 4**. Chloroplasts in bundle sheath of 10103 $T_2$ homozygous lines have normal ultrastructure. Chloroplasts in mestome sheath of nulls and 10103 $T_2$ homozygous lines showed differences in chloroplast development and ultrastructure. WT Iis Kitaake; OE-ZmG2-E131 is G2 OE; 6-15 is G2&PDVs Null 1, 6-5 is G2&PDVs OE 1 ; 20-8 is G2&PDVs Null 2, 20-5 is G2&PDVs 2; 25-8 is G2&PDVs Null 3, 25-7 is G2&PDVs OE 3.

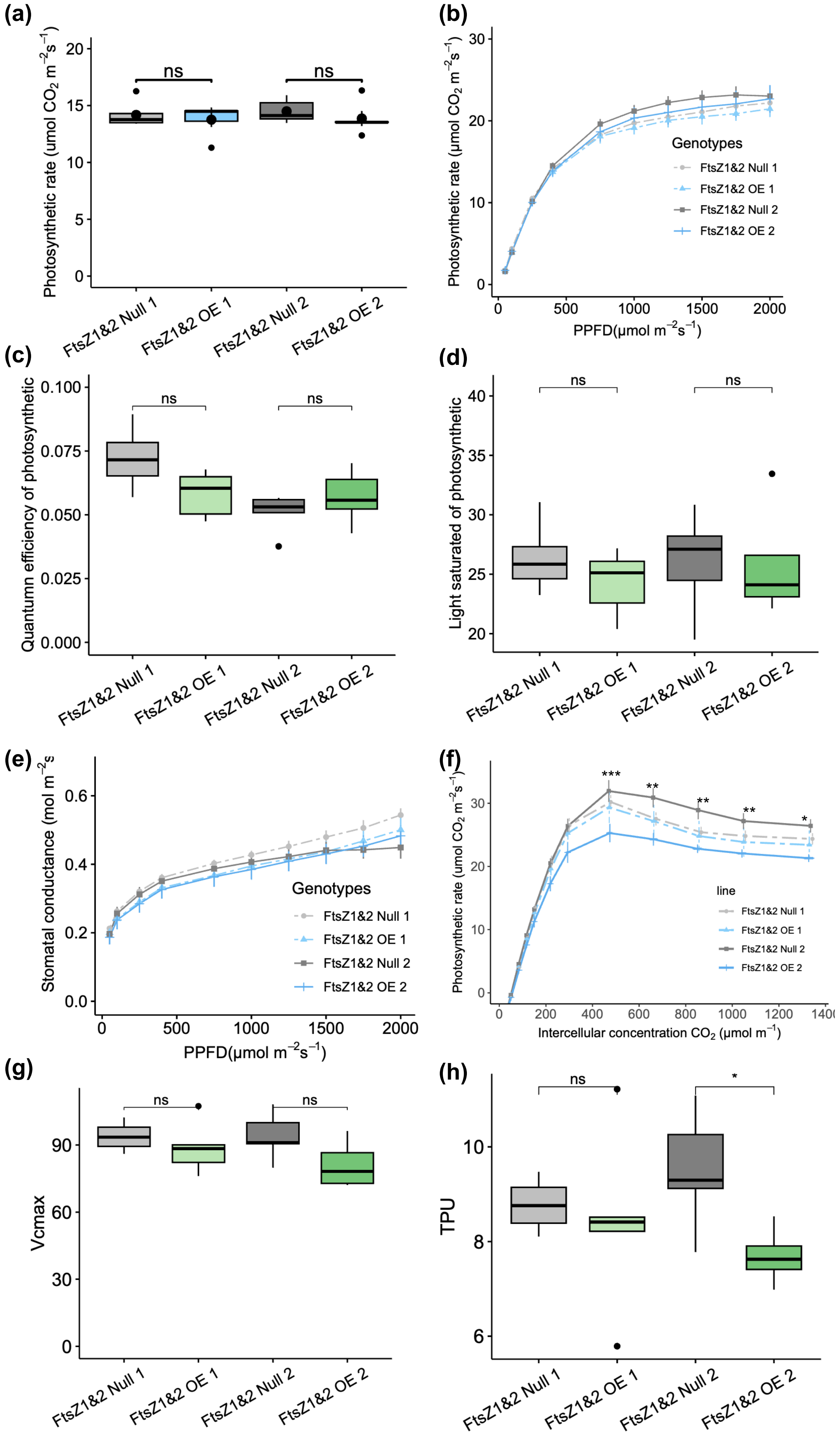


**Fig. S10 Photosynthesis rates declined under high $CO_2$ in the strong overexpression lines of *OsFtsZ1&2* (a)** Net photosynthetic rate under ambient conditions (400 ppm $CO_2$, 400 μmol $m^{-2}$ $s^{-1}$ PPFD). **(b)** Light response curve of net photosynthesis (n = 6); (**c**) The Quantum yield of photosynthetic; (**d**) The light saturated of photosynthetic; **(e)** Stomatal conductance (n=6). **(f)** A–Ci response curve (n=6). Line *Ubi$_{pro}$:FtsZ1&2* OE 2, with high expression of *OsFtsZ1&2*, exhibited a reduced photosynthetic rate at intercellular $CO_2$ concentrations above 400 ppm. Data represent means ± SD from six independent biological replicates per line. (**g-h)** Vcmax (g) and TPU (h) generated by A-Ci curve.

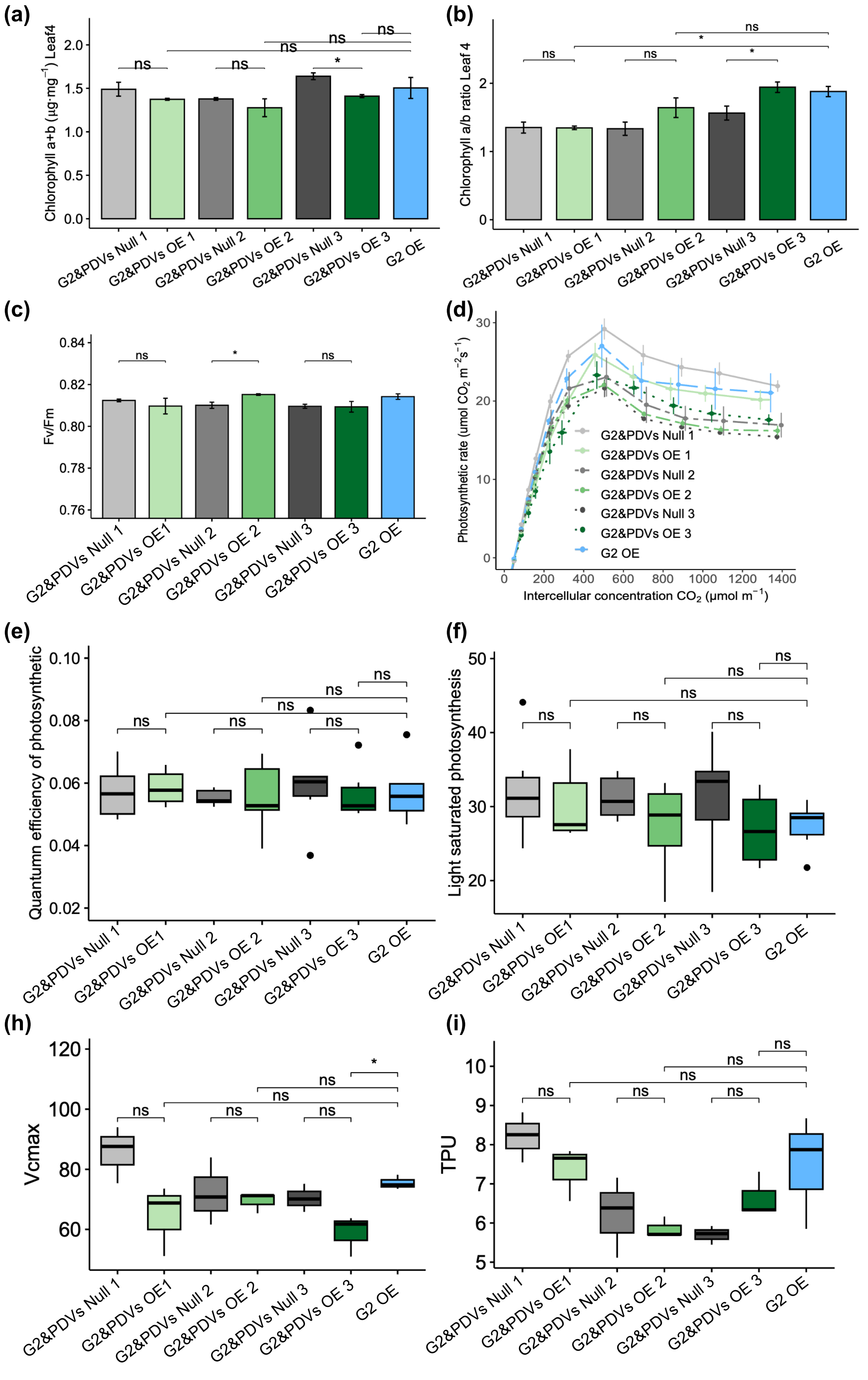


**Fig. S11 Photosynthesis parameters are not altered in rice lines expressing *ZmG2, OsPDV1&2.*** (a, b) Total chlorophyll (a) and chlorophyll a/b ratio (b); (c) maximal photosynthetic efficiency of photosystems II (Fv/Fm); (d) A-Ci response curve; (f) The Quantum yield of photosynthetic; (g) The light saturated of photosynthetic; (h-i) Vcmax (h) and TPU (i) generated by A-Ci curve

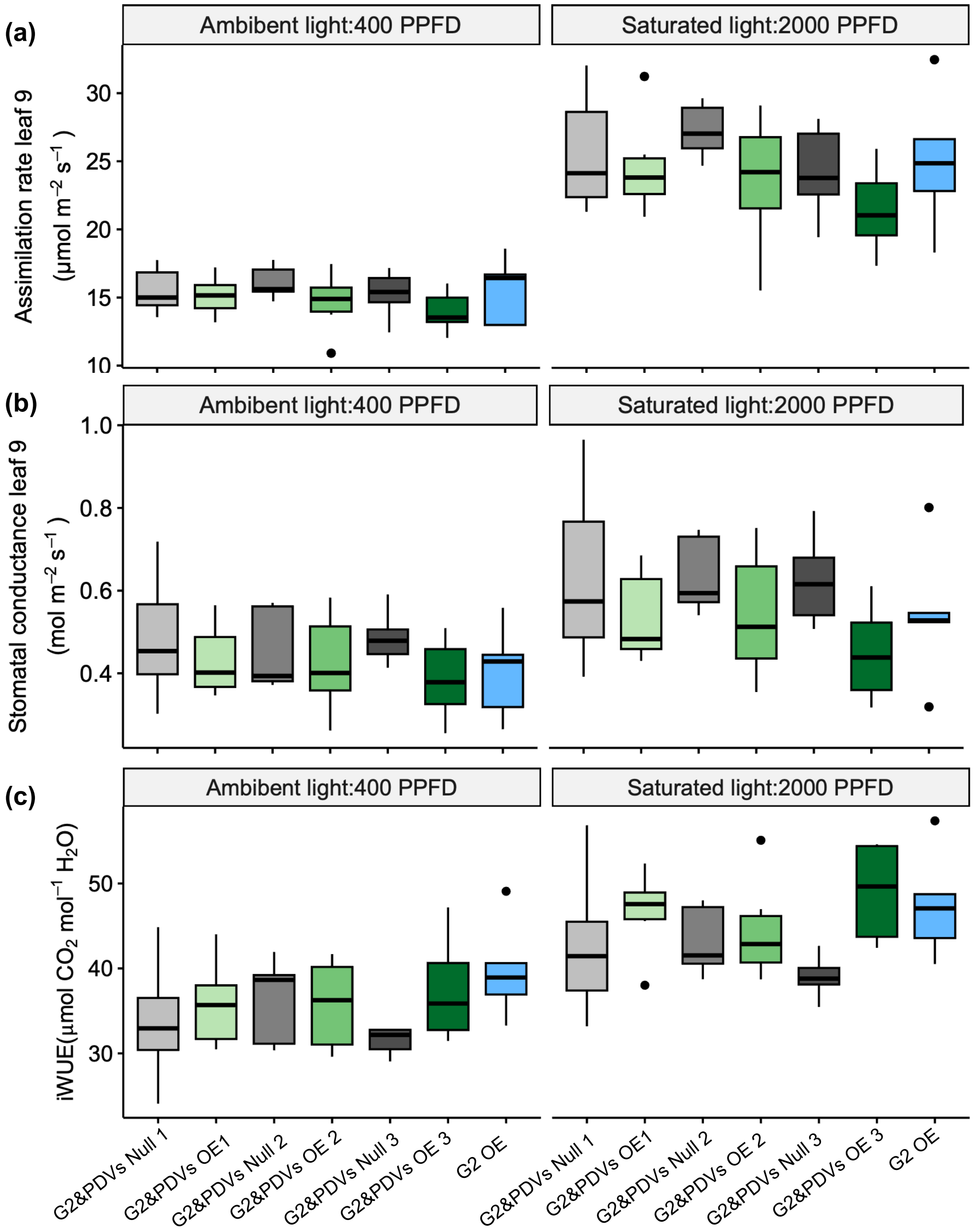


**Fig. S12 Overexpression of *ZmG2, OsPDV1* and *OsPDV2* showed high intrinsic water use efficiency.** (a-c) Assimilation rate (a), stomatal conductance (b) and intrinsic water use efficiency (c) of the 9th leaf under ambient (400 PPFD) and saturated (2,000 PPFD) light conditions after a 10-min stabilization.

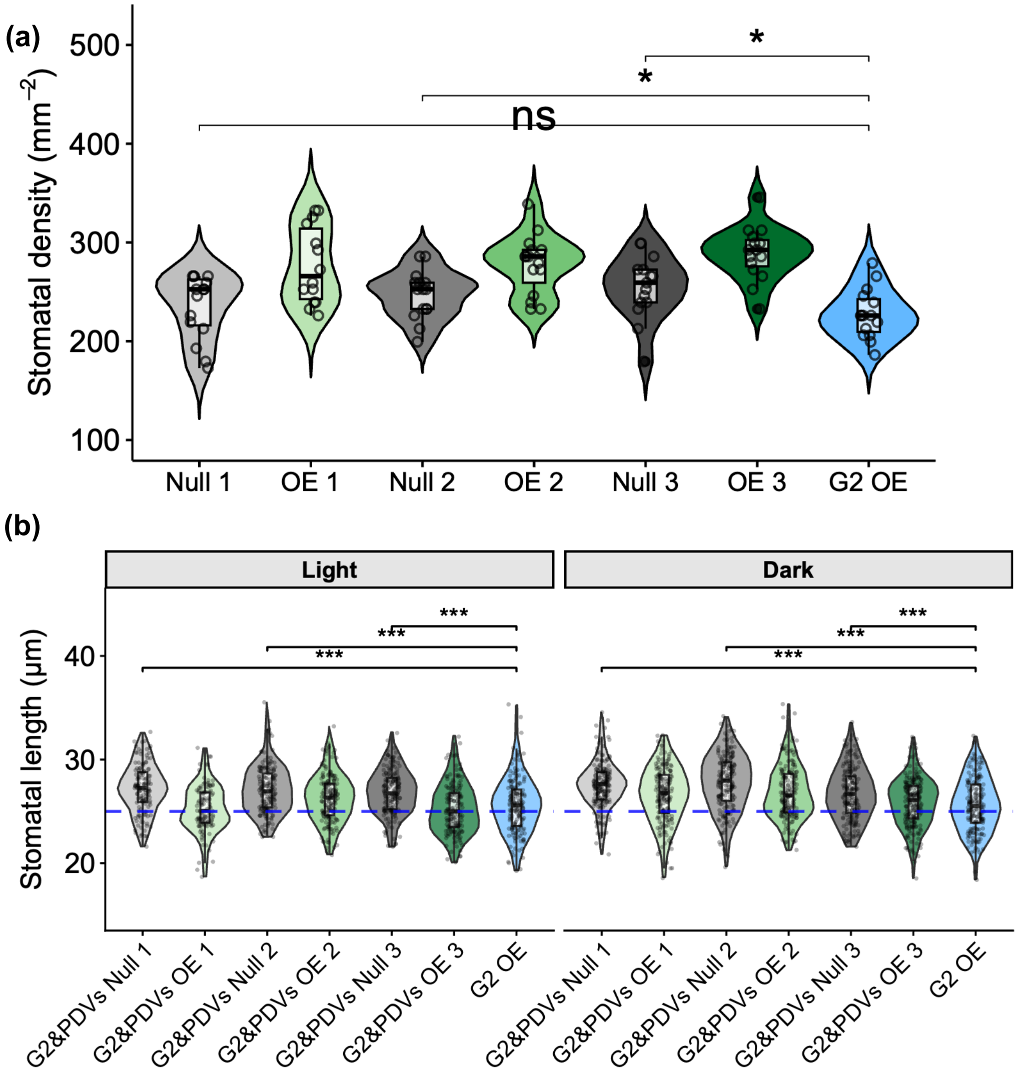


**Fig. S13 Overexpression of *ZmG2* reduced the stomatal density and stomatal length compared to nulls.** (a) Quantification of stomatal density, showing statistically significant differences between *OE-ZmG2* compared with null. (b) Stomatal length under light and dark conditions on the abaxial leaf blade showing reductions in overexpression *ZmG2* lines. Stars above violin plots indicate statistically significant differences between *ZmG2* and nulls, as determined by independent *t*-test: $p \leq 0.05$ (*), $p \leq 0.01$ (**), $p \leq 0.001$ (***). Non-significant comparisons are labelled as “ns”.
